# Inhibition of RNA splicing is a novel therapeutic strategy for disruption of nuclear replicating viruses

**DOI:** 10.64898/2026.06.26.734849

**Authors:** Lorenzo Serra, Claire M. O’Brien, Molly R. Patterson, Clayton J. Otter, Alison Yu, Reyes W. Acosta, Daniel T. Claiborne, Susan R. Weiss, Alexander M. Price

**Affiliations:** Genome Regulation and Cell Signaling, Ellen and Ronald Caplan Cancer Center, The Wistar Institute, Philadelphia, PA, United States; Cellular and Molecular Biology Program, Department of Pharmacy and Biotechnology, University of Bologna, Bologna, Italy; Department of Microbiology, Perelman School of Medicine, University of Pennsylvania, Philadelphia, PA, United States; Cell and Molecular Biology Graduate Group, Perelman School of Medicine, University of Pennsylvania, Philadelphia, PA, United States; Penn Center for Research on Emerging Viruses, Perelman School of Medicine, University of Pennsylvania, Philadelphia, PA, Unites States

## Abstract

RNA splicing is a fundamental feature of eukaryotic gene expression that is co-opted by many nuclear-replicating viruses to produce viral transcripts. Recent clinical development of spliceosome-targeting therapeutics has demonstrated that splicing can be safely modulated *in vivo,* raising the possibility that viral dependence on host RNA processing may represent an exploitable antiviral vulnerability. Here, we show that pharmacologic inhibition of cellular RNA splicing preferentially suppresses the expression of spliced viral transcripts and profoundly impairs replication of multiple splicing-dependent viruses. In human adenovirus, low-dose inhibition of the spliceosome using mechanistically distinct chemical inhibitors, or genetic depletion of SF3B1, disrupted splicing of complex late viral transcripts while largely sparing simple early transcripts. This resulted in impaired viral DNA replication, reduced late protein accumulation, and multi-log decreases in infectious progeny production. Splicing inhibitors similarly impaired RNA splicing and viral replication of DNA virus Herpes Simplex virus 1 and RNA virus influenza A, which both rely on host-dependent RNA processing strategies. In contrast, cytoplasmic RNA viruses lacking spliceosome dependence were unaffected, supporting an on-target mechanism. Together, these findings identify viral RNA processing complexity as a determinant of antiviral sensitivity and establish host spliceosome dependence as a shared and targetable vulnerability across diverse viral families.

## Introduction

RNA processing is a fundamental feature of eukaryotic cells that coordinates gene expression and expands coding capacity through proteomic diversity^1^. In addition, most nuclear-replicating viruses require cellular RNA Polymerase II and/or associated splicing machinery for the expression of viral genes^2^. Therefore, viral infection often creates a state in which cellular RNA processing machinery becomes extensively co-opted for viral replication^3^. Such a strong dependency on a host process opens the possibility for broad-spectrum host targeting antiviral drugs, but the essentiality of RNA processing for cell viability has raised substantial concerns about the toxicity of any such pharmacologic inhibition. Nevertheless, specific inhibitors of eukaryotic RNA splicing have recently entered human clinical trials for the treatment of cancers and other malignancies driven by spliceosome mutations^4^. It is unknown whether viral RNA transcripts are uniquely sensitive to the disruption of cellular splicing.

Human adenoviruses (AdV) provide an especially powerful system for understanding the dynamic relationship between viral replication and cellular RNA processing^5^. Indeed, pioneering studies of AdV led to the discovery of cellular RNA splicing itself^6,7^. AdV gene expression is entirely dependent on cellular transcriptional machinery, yet the extent of RNA processing for viral transcripts differs dramatically over the course of infection^8^. Early transcripts are generally derived from individual transcriptional units with relatively limited alternative processing. In contrast, the major late transcriptional unit generates more than 40 mRNA isoforms from a single primary transcript through extensive alternative splicing and polyadenylation. This transition from relatively simple early transcription to complex late-stage RNA processing suggests that distinct stages of the viral lifecycle may preferentially utilize key aspects of host splicing machinery. Previous studies have largely focused on individual RNA binding proteins or isolated viral transcripts rather than systematic perturbation of spliceosome function itself.

In addition, diverse viruses utilize a breadth of RNA splicing mechanisms. Like AdV, Herpes Simplex viruses (HSV) are double-stranded (ds)DNA viruses that replicate in the nucleus and exclusively use cellular RNA polymerases to drive viral gene expression^9^. However, while HSV-1 encodes over 100 distinct transcripts, most herpes transcripts are intronless with only five routinely spliced by cellular machinery^10,11^. Although most of these spliced HSV-1 transcripts are dispensable for the lytic cycle of infection, they nevertheless encode critical functional regulators of viral infection. On the opposite end of the spectrum, influenza viruses have single stranded negative sense segmented RNA genomes that replicate within the nucleus. While Influenza viruses utilize a viral RNA-dependent RNA polymerase for mRNA transcription, two key segments, M and NS, are subsequently spliced into essential alternative mRNA isoforms^12^. These RNAs are spliced post-transcriptionally, but still require host splicing machinery to coordinate this unconventional splicing reaction^13,14^. These RNA processing strategies suggest that different viruses may exhibit differential sensitivity to drugs targeting distinct stages of the splicing cycle.

Recent advances in spliceosome-targeting therapeutics have enabled pharmacologic interrogation of these questions^4^. Small molecules including pladienolides, spliceostatins, and herboxidienes impair spliceosome assembly through interactions with components of the Splicing Factor 3B (SF3B) complex, a critical mediator of branchpoint recognition during spliceosome formation^15–17^. Several of these compounds, including the pladienolide derivative E7107 and the orally bioavailable H3B-8800, have advanced into clinical trial for spliceosome-mutant malignancies such as Acute Myelogenous Leukemia and Myelodysplastic Syndrome^18,19^. Importantly, studies with *in vitro* cancer systems have demonstrated that chemical inhibition of splicing does not affect all transcripts equally, but instead preferentially impacts RNAs with weak splice sites, short intronic sequences, or higher than expected GC content^20^. In addition, inhibition of certain cyclin-dependent kinases can uncouple transcript elongation from co-transcriptional splicing^21,22^. These findings raise the possibility that clinically achievable modulations of cellular RNA splicing could be repurposed to target selective vulnerabilities in the aforementioned nuclear replicating viruses and their critical need to splice viral transcripts.

Here we demonstrate that inhibition of cellular RNA splicing preferentially suppresses the expression of complex viral transcripts and profoundly impairs replication of multiple splicing-dependent viruses. Using pharmacologic inhibitors of spliceosome function, as well as genetic depletion of SF3B1, we find that adenoviral late transcripts exhibit substantially greater sensitivity to impaired splicing than early viral transcripts. Blocking splicing of late viral transcripts leads to significant defects in viral genome replication, late protein accumulation, and infectious progeny production. This vulnerability extends to primary cell models of infection and evolutionarily distinct viruses that utilize host-mediated RNA splicing, including HSV-1 and multiple Influenza A viruses. Importantly, cytoplasmic RNA viruses that do not employ spliceosomal processing are not affected by the same treatments, implicating direct effects via splicing modulation and not pleiotropic toxicities. Together, these findings identify alternative viral RNA processing as a determinant of antiviral sensitivity, establishing host spliceosome dependence as a targetable vulnerability across diverse virus families.

## Results

### Adenovirus infection is suppressed by pladienolide splicing inhibition

SF3B1-targeting pladienolides are the most well-characterized of the chemical splicing inhibitors, so we began our studies by focusing on Pladienolide B and the second generation, clinical-grade compound H3B-8800. We hypothesized that adenoviruses (AdV), the family of viruses where RNA splicing itself was discovered, would be particularly sensitive to inhibition of cellular splicing. A549 lung adenocarcinoma cells were infected with a high multiplicity of infection (MOI 10) of adenovirus serotype 5 (Ad5). Two hours post infection (hpi) we treated cells with a wide dose range of both Pladienolide B (PladB) and H3B-8800 (H3B) and assessed adenoviral early and late protein expression by western blot (**Figure 1A-B**). The chosen concentrations spanned the high nanomolar to low micromolar doses previously shown to be effective against spliceosome-mutated cancer cells^15,20^. While all tested concentrations had modest effects on early viral proteins E1A and DBP, the decrease in late viral proteins was substantially more apparent. This differential effect in late protein and early protein accumulation reached maximum disparity at approximately 1 nM for PladB and 30 nM for H3B. Since viral late gene expression is dependent upon successful viral early gene expression and DNA replication, we treated Ad5-infected A549 cells at the very beginning of the late stage (16 hpi) (**Figure 1C**). At this time point early viral protein production (E1A, DBP) was not appreciably affected, however viral late product accumulation was still depleted compared to the vehicle control. These findings imply that relatively low doses of these splicing inhibitors have preferential effects on the production of the late class of adenovirus protein production.

**Figure 1.**
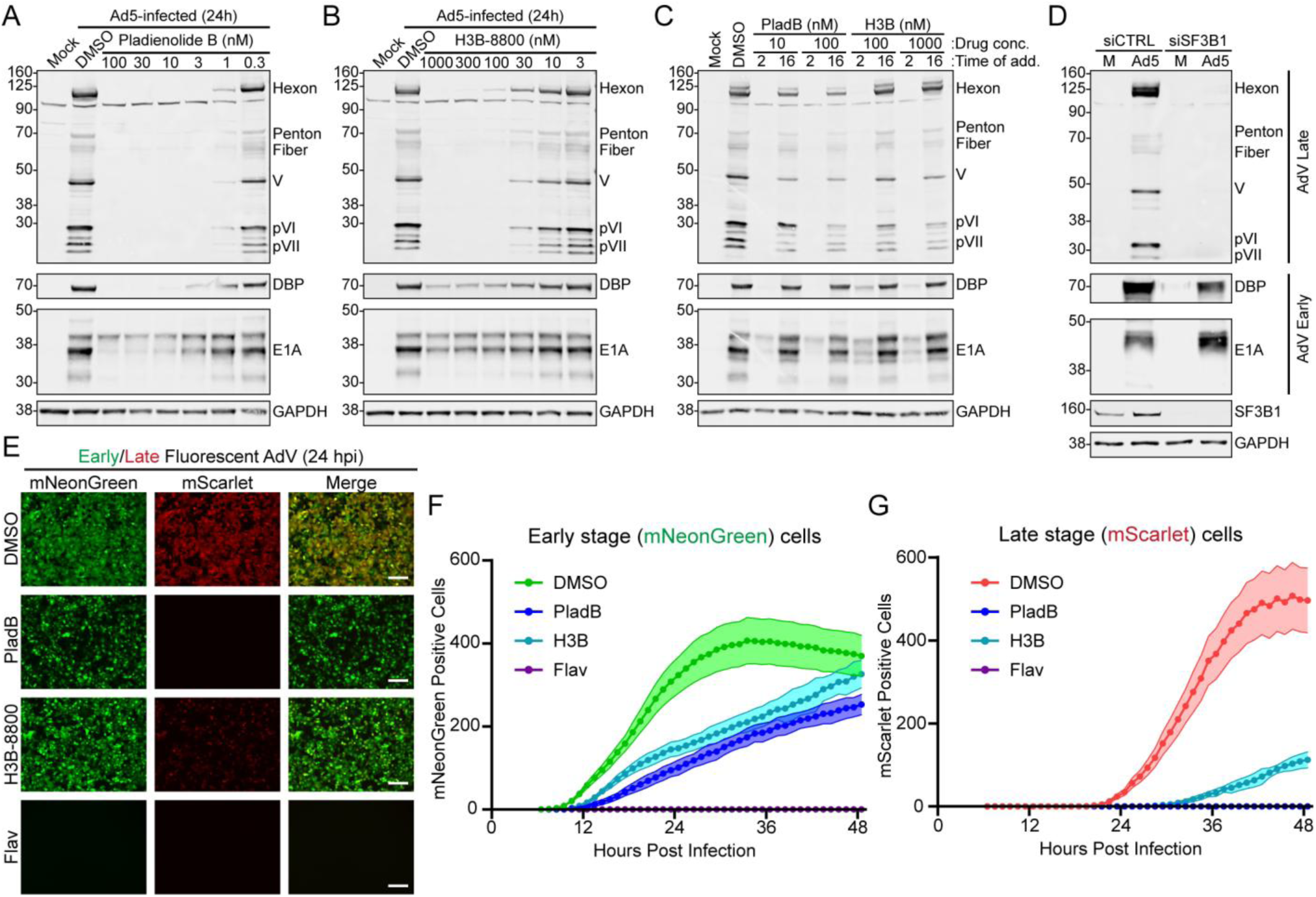
Splicing inhibition displays a stage-dependent effect on adenovirus gene expression. **(A)** Immunoblot of uninfected A549 cells (Mock) or Ad5-infected A549 cells treated with vehicle control or different concentrations of PladB at 2 hpi or **(B)** H3B-8800. Lysates were collected at 24 hpi and probed for Ad5 capsid proteins to assess late expression and DBP and E1A to assess early expression. GAPDH was used as a loading control. **(C)** A549 cells were treated with either low or high concentrations of PladB or H3B at 2 hpi or 16 hpi. Lysates were collected at 24 hpi and probed for Ad5 early and late proteins. **(D)** A549 cells were transfected with a pool of siRNA targeting SF3B1 for 24 hours. Cells were subsequently mock infected (M) or Ad5-infected for an additional 24 hours before lysates were collected for early and late protein immunoblot. **(E)** A549 cells were infected with a dual-fluorescence reporter of Ad5 infection (E3gL5s) and treated with vehicle DMSO, 100 nM PladB, 1000 nM H3B, or 300 nM flavopiridol (Flav) at 2 hpi. At 24 hpi cells were imaged for early green mNeonGreen or late red mScarlet3 fluorescence. White scale bar represents 100 μm. **(F)** CellCyte analysis of early reporter green fluorescence positive A549 cells following E3gL5s infection imaged every hour for 48 hpi. DMSO (green), 100 nM PladB (blue), 1000 nM H3B (teal), or Flav (purple) were added at 2 hpi. **(G)** CellCyte analysis of late reporter red fluorescence positive A549 cells following E3gL5s infection imaged every hour for 48 hpi. DMSO (red), 100 nM PladB (blue), 1000 nM H3B (teal), or 300 nM Flav (purple) were added at 2 hpi. For F-G the shaded area represents the SEM of six replicates.

Both PladB and H3B target the SF3B1 subunit that recognizes branchpoint sites in pre-mRNA splicing. To determine if genetic depletion of SF3B1 phenocopies the results of the small molecule inhibition, we transfected A549 cells with control or SF3B1-targeting siRNA 24 hours before infection. Subsequently, cells were infected with Ad5 and harvested 24 hours post infection for a total of 48 hours of siRNA-mediated knockdown before western immunoblot for early and late viral proteins (**Figure 1D**). SF3B1 knockdown was confirmed to be efficient in these conditions. Similar to the effects of chemical inhibition, SF3B1-depletion led to no appreciable effect on viral early protein accumulation but a profound effect on viral late protein accumulation. These results confirm that genetic disruption of RNA processing has a substantial and preferential effect on adenovirus late protein production.

Our laboratory recently developed a fluorescent reporter virus for monitoring early- and late-stage effects during adenovirus infection^23^. In this system, early gene production is measured by green fluorescence from mNeonGreen expressed from the native E3 gene locus while late gene production is measured by red fluorescence from mScarlet3 cleaved from a P2A site from the c-terminus of the late Fiber gene. At 24 hpi, vehicle-treated cells displayed both green and red fluorescence, indicating successful early and late gene production. However, treatment with either PladB or H3B entirely abrogated the late red fluorescence while sparing early green fluorescence (**Figure 1E**). This effect was different from treatment with a broad-spectrum inhibitor of RNA Pol II transcription, the CDK9-inhibitor flavopiridol, which was able to block both early and late reporter fluorescent expression. Additionally, early green fluorescence (**Figure 1F**) or late red fluorescence (**Figure 1G**) was tracked in living cells in real time using a CellCyte system. Early gene reporter expression was delayed in accumulation after treatment with PladB or H3B compared to the vehicle control, yet at the end of the 48 hour experiment all treatments had the same level of cells positive for infection. In contrast, late gene expression was entirely blocked by PladB or Flavopiridol treatment, and substantially attenuated by H3B treatment. Together, these results demonstrate that splicing inhibition slows the progression of early gene accumulation as well as substantially blocks the switch to late-stage infection and ultimately the accumulation of late-stage viral proteins.

### Splicing inhibition has a preferential effect on the splicing of late viral genes

All adenovirus genes are processed and spliced by cellular machinery, and this splicing is thought to take place co-transcriptionally with RNA Pol II. Viral early genes are considered “simple”, with generally one promoter site, limited alternative cleavage and polyadenylation, and limited internal splicing events. In contrast, late transcripts are “complex” and transcribed from only a single genetic unit yet alternatively spliced and polyadenylated to create over 40 unique RNA isoforms (**Supplementary Figure 1A**). We hypothesized that this extreme reliance on alternative RNA processing could explain the apparent sensitivity of late gene expression to splicing inhibitors.

We proceeded to determine the effect of PladB and H3B treatment on the accumulation of specific viral transcripts. We tested diverse early transcripts (E1A, E2A-DBP, and common E4) as well as late transcripts (Spliced tripartite leader (TPL), L2-V, and L5-Fiber). A549 cells were infected with Ad5 at MOI 10, treated with drugs at 2 hpi, and RNA was collected for quantitative reverse transcriptase PCR at 24 hpi (**Figure 2A**). While the differential accumulation of early transcripts was varied, many transcripts were not significantly different from vehicle control or less than two-fold decreased. In contrast, all tested late transcripts were significantly diminished. Splice-specific qPCR was used to interrogate this finding on select early- and late-gene splicing events further (**Supplementary Figure 1A-B**). While E1A was unaffected by treatment with PladB or H3B, the spliced isoforms of either late TPL or Fiber were significantly decreased, while the unspliced pre-mRNA forms were less affected. This led to a significant decrease in the efficiency of splicing for late genes but not the early gene, defined as the ratio of spliced to unspliced specific transcript (**Figure 2B**). These results show that low doses of splicing inhibitors have preferential effects on the alternative splicing of late viral genes, but more muted effects on the essentially constitutive splicing of early viral genes.

**Figure 2.**
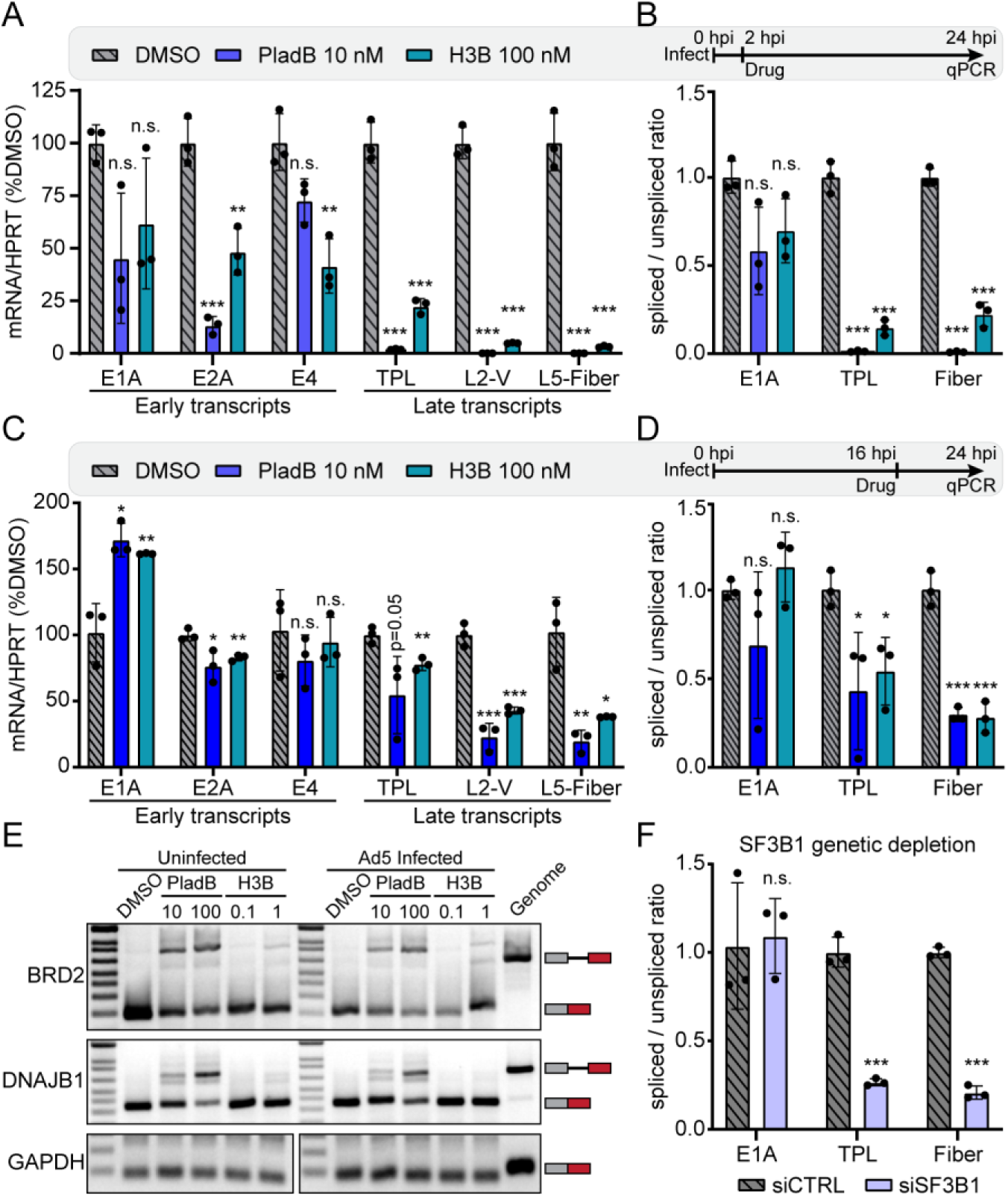
Splicing inhibition predominantly impacts complex late viral gene splicing. **(A)** A549 cells were infected with Ad5, then treated with the specified concentrations of PladB (blue) or H3B (teal) at 2 hpi. Total RNA was harvested at 24 hpi and qRT-PCR performed for listed viral early and late transcripts and normalized to both cellular HPRT1 transcripts and DMSO-treated vehicle controls. Data points are shown for three biological replicates, bars depict mean, and error bars show standard deviation. **(B)** The unspliced transcript levels from A were calculated by qRT-PCR and the ratio of spliced to unspliced transcript was plotted. Data points are shown for three biological replicates, bars depict mean, and error bars show standard deviation. **(C)** Same as in A, but drugs were added at 16 hpi before RNA was harvested and analyzed by qRT-PCR at 24 hpi. **(D)** Same as in B, but the splicing ratio was calculated from the 16-24 hpi treatments shown in C. **(E)** RT-PCR agarose gel showing intron retention between exons 4-5 of human BRD2 and exons 2-3 of human DNAJB1. Control exon PCR is shown for GAPDH as a loading control. All A549 cells were treated with the listed concentrations of PladB or H3B for eight hours. In the Ad5-infected conditions the cells were virus infected for 16 hours before the 8 hour drug treatment. A549 genomic DNA was substituted for reverse transcribed cDNA as a positive unspliced control. **(F)** A549 cells were transfected with a pool of siRNA targeting SF3B1 for 24 hours. Cells were subsequently Ad5-infected for an additional 24 hours before total RNA was collected and qRT-PCR performed for viral early and late spliced and unspliced transcripts. Data points are shown for three biological replicates, bars depict mean, and error bars show standard deviation. For all data, significance was analyzed by unpaired two-tailed *t*-test. Significance was shown as P-value >0.5 (not-significant, n.s.), * P< 0.05, ** P< 0.01 and *** P< 0.001.

To disentangle if the modest effects on early gene transcription were ultimately responsible for the observed strong defects in late gene expression we waited until 16 hours post infection before treating with either low or high doses of both PladB and H3B and analyzing RNA expression 8 hours later at 24 hpi. Similar to above, low doses of PladB and H3B had no effect, subtle decreases, or even subtle increases on early gene expression (**Figure 2C; Supplementary Figure 2C**). Once again, the effects on late gene expression, especially the more distal L2-V and L5-Fiber, were strongly decreased. This effect included a specific decrease in late gene splicing efficiency (**Figure 2D**). In this shorter time scale experiment, higher doses of both drugs (100 nM PladB, 1 μM H3B) had minimal effects on early genes, but significant effects on late gene accumulation and splicing (**Supplementary Figure D-E**). To validate that this limited window of treatment affected cellular transcript splicing, we performed reverse transcriptase PCR on two cellular gene intron retention events known to be sensitive to these drugs, BRD2 and DNAJB1^15^ (**Figure 2E**). Both PladB and H3B caused a dose-dependent increase in cellular intron retention. Importantly, concurrent adenovirus infection did not alter these effects on cellular splicing. These results confirm that splicing inhibitors impair the splicing efficiency of late viral transcripts.

To determine if genetic depletion of SF3B1 recapitulates the effect of chemical inhibition on pre-mRNA splicing, we depleted SF3B1 by siRNA 24 hours before infection and subsequently collected RNA 24 hpi (**Figure 2F**). SF3B1 knockdown had no significant effect on the splicing efficiency of early E1A, but substantially decreased splicing of late TPL and Fiber. Together, these results indicate that viral late transcripts with more complicated splicing patterns are more sensitive to changes in cellular RNA processing compared to viral early genes.

### Splicing inhibitors profoundly suppress adenoviral replication and progeny production

Given the effect of splicing inhibitors on both adenovirus transcription and protein production, we hypothesized that these drugs would also impair viral replication. We began by measuring viral DNA replication inside infected A549 cells after treating with a dose curve of both PladB and H3B at 2 hpi (**Figure 3A-B**). At all times post-infection viral DNA accumulation was impaired compared to vehicle control; however, the magnitude of this effect waned as the infection progressed or was not significant at the lowest dose measured. This finding was partially surprising, as DNA replication is mediated by the early viral proteins that were less affected by these drugs at low concentrations. To determine if the modest effects seen on early viral proteins at the 2 hpi dosing time mediated this effect on viral DNA replication, we infected A549 cells and subsequently treated them at 16 hpi, a time that was previously shown not to affect early protein accumulation (**Figure 1C**). In this late treatment window, neither splicing inhibitor had a significant effect on viral DNA replication (**Figure 3C**). These data imply that even modestly impacting early viral proteins via splicing inhibition can delay the process of viral replication and impair accumulation of viral DNA.

**Figure 3.**
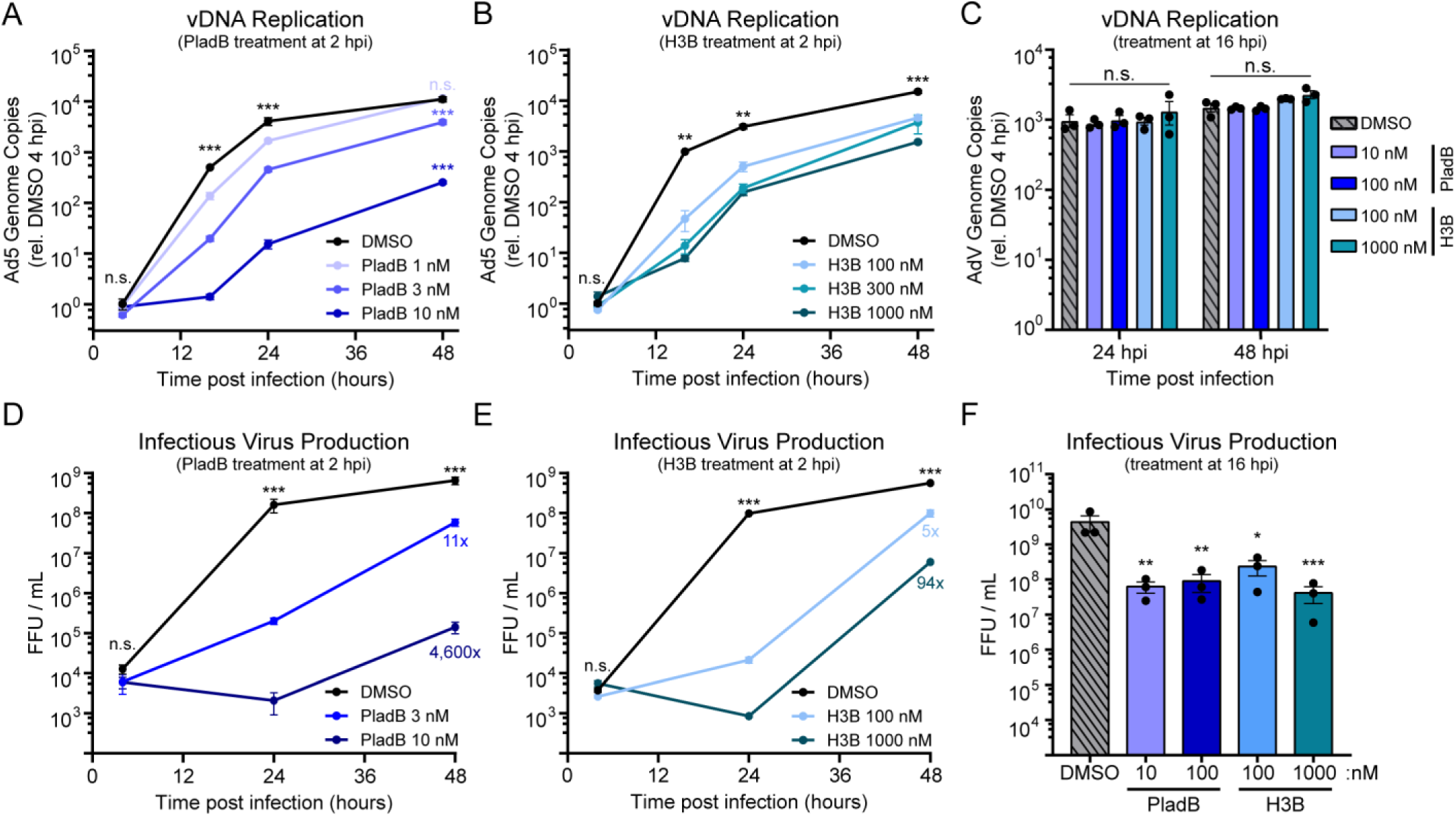
Splicing inhibition strongly impairs viral replication and infectious progeny production. **(A)** A549 cells were infected with Ad5 and then treated at 2 hpi with the listed concentrations of PladB. At the listed times DNA was harvested and subjected to qPCR with primers targeting an Ad5 genomic region and normalized to both cellular tubulin DNA and relative viral DNA was normalized to cellular tubulin and input viral DNA at 4 hpi. Data are presented as log-scale, points show the average of three biological replicates, and error bars show standard deviation. **(B)** Same as A, but A549 cells were treated with the listed concentrations of H3B. **(C)** A549 cells were infected with Ad5 with subsequent treatments of low or high dose PladB (blue) or H3B (teal) added at 16 hpi. DNA was collected at 4, 24, or 48 hpi and relative viral DNA was normalized to cellular tubulin and input viral DNA at 4 hpi. **(D)** A549 cells were infected with fluorescent Ad5 (E3mNG) and treated with listed concentrations of PladB at 2 hpi. Total cell plus supernatant was collected at 4, 24, or 48 hpi and released viruses used for virus progeny calculation via fluorescent forming unit (FFU). Data are presented as log-scale, points show the average of three biological replicates, and error bars show standard deviation. **(E)** Same as D, but A549 cells were treated with the listed concentrations of H3B. **(F)** A549 cells were infected with fluorescent Ad5 and subsequently treated with low or high dose PladB (blue) or H3B (teal) at 16 hpi. Total cell plus supernatant was collected at 48 hpi and released viruses used for virus progeny calculation via fluorescent forming unit (FFU). For all statistical analyses significance was analyzed by lognormal *t*-test. Significance was shown as P-value >0.5 (not-significant, n.s.), * P< 0.05, ** P< 0.01 and *** P< 0.001.

We next looked at the ability of splicing inhibitors to impair the production of fully infectious virus by collecting intact cells over a time course and performing fluorescence focus forming assays. At both low and high doses of PladB and H3B the production of infectious virus was impaired by multiple logs (**Figure 3D-E**). Since the production of mature viral particles is the ultimate step in the viral lifecycle any defect in the prior steps, such as DNA replication, could have substantial knock-on effects. To rule out the substantial loss of infectious particles being solely mediated by poor DNA replication, we treated infected A549 cells at 16 hpi, a time that led to no detectable changes in viral DNA accumulation, and subsequently assayed for infectious virus at 48 hpi (**Figure 3F**). Under this dosing strategy, infectious virus production was still impacted by 1-2 logs (21x-130x fold depletion). Taken together, these results show that even in the absence of an effect on viral DNA replication, late treatment can still strongly affect splicing of viral late transcripts, leading to a substantial decrease in viral infection.

### Alternative splicing inhibitors that act through different mechanisms impair viral replication

Both PladB and H3B are pladienolide compounds that act by binding SF3B1 to impair branch point adenosine recognition in the A Complex of the spliceosome^15,20^ (**Figure 4A**). Alternative compounds have been described and characterized to inhibit pre-mRNA processing, including Isoginkgetin (IsoG)^24^, a biflavinoid compound with unknown specific mechanism of action that stalls the U4/U5/U6 tri-SNRNP complex from forming the B Complex of the spliceosome, and OTS-964 (OTS)^21,25^, a CDK11 inhibitor that has been shown to block the kinase involved in SF3B1 phosphorylation, a necessary step for the release of the branch-site adenosine during the activated B* Complex of the spliceosome. Considering the less well characterized role of these compounds on splicing, we confirmed their effect on human genes by RT-PCR in A549 cells with two concentrations in the range that was described to produce an effect on splicing in the literature (**Figure 4B**). We hypothesized that despite their different mechanisms of action they would still inhibit adenovirus infection in a splicing-dependent manner.

**Figure 4.**
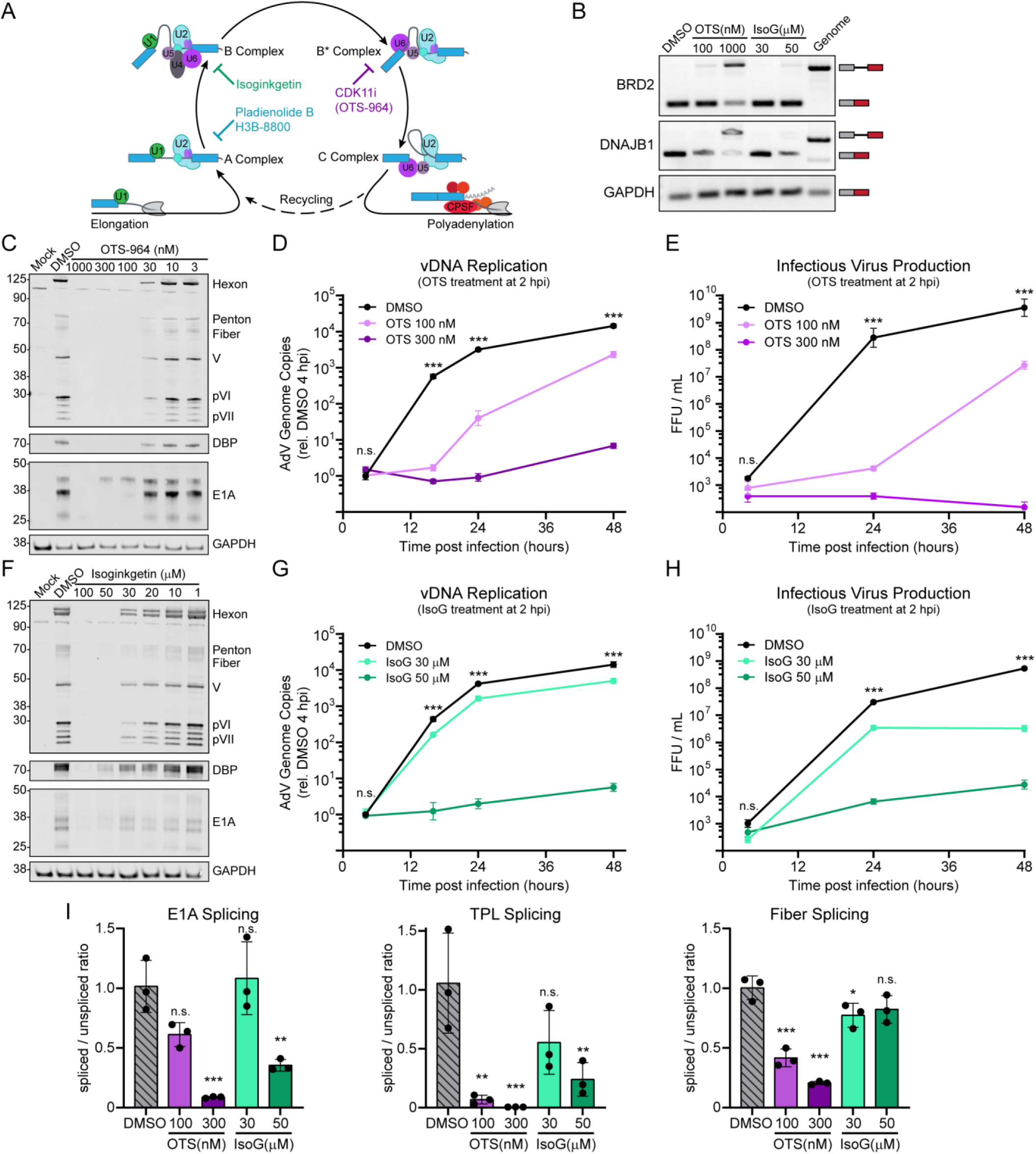
Non-pladienolide splicing inhibitors similarly affect viral replication. **(A)** Schematic diagram showing the stepwise recruitment of U1, U2, and U4/5/6 snRNPs to form the distinct complexes of the splicing cycle on a pre-mRNA. Steps that are specifically blocked by PladB or H3B, Isoginkgetin (IsoG), or OTS-964 (OTS) are indicated. **(B)** RT-PCR agarose gel showing intron retention between exons 4-5 of human BRD2 and exons 2-3 of human DNAJB1. Control exon PCR is shown for GAPDH as a loading control. All A549 cells were treated with the listed concentrations of OTS or IsoG for eight hours. A549 genomic DNA was substituted for reverse transcribed cDNA as a positive unspliced control. **(C)** Immunoblot of uninfected A549 cells (Mock) or Ad5-infected A549 cells treated with vehicle control or different concentrations of OTS at 2 hpi. Lysates were collected at 24 hpi and probed for Ad5 early and late proteins. GAPDH was used as a loading control. **(D)** A549 cells were infected with Ad5 and then treated at 2 hpi with the listed concentrations of OTS. DNA was collected at listed times and relative viral DNA was normalized to cellular tubulin and input viral DNA at 4 hpi. **(E)** A549 cells were infected with fluorescent Ad5 (E3mNG) and treated with listed concentrations of OTS at 2 hpi. Total cell plus supernatant was collected at listed times and released viruses used for virus progeny calculation via fluorescent forming unit (FFU). Data are presented as log-scale, points show the average of three biological replicates, and error bars show standard deviation. **(F)** Same as in C for A549 cells treated with IsoG. **(G)** Same as in D for A549 cells treated with IsoG. **(H)** Same as in E for A549 cells treated with IsoG. **(I)** A549 cells were infected with Ad5 and subsequently treated with low or high doses of OTS or IsoG at 16 hpi. Total RNA was collected and qRT-PCR performed for viral early and late spliced and unspliced transcripts and splicing ratio was plotted. Data points are shown for three biological replicates, bars depict mean, and error bars show standard deviation. For qRT-PCR significance was analyzed by unpaired two-tailed *t*-test. For vDNA and FFU quantification significance was analyzed by lognormal *t*-test. Significance was shown as P-value >0.5 (not-significant, n.s.), * P< 0.05, ** P< 0.01 and *** P< 0.001.

A549 cells were first infected with high MOI 10 of Ad5 before being treated with a wide dose range of both OTS and IsoG at 2 hpi. Infected cells were collected at 24 hpi and immunoblotted for viral early and late proteins (**Figure 4C,F**). Similar to PladB and H3B, these alternative splicing inhibitors led to a stronger defect in late protein accumulation compared to early protein accumulation, with the inflection points of highest disparity at 100 nM for OTS and 30 μM for IsoG. We then ascertained the effects of these two drugs on viral DNA replication by performing qPCR over a time course of infection after early treatment with high- and low-doses of both drugs at 2 hpi (**Figure 4D,G**). OTS showed very strong efficacy in repressing viral DNA replication with log-scale reductions at all tested time points for both 300 nM and 100 nM doses. IsoG, on the other hand, entirely blocked viral replication at the high dose of 50 μM, but showed more modest yet significant effects at 30 μM. This dose-dependent effect of IsoG mirrors the effect on host gene splicing (**Figure 4B**). Finally, infectious progeny production was measured by fluorescent focus forming assay at 24 and 48 hpi after an early treatment with the same doses at 2 hpi (**Figure 4E,H**). Log-scale decreases in infectious virus production were observed for all drugs and dosages at each timepoint. These data demonstrate that drugs known to impair alternative steps of the cellular splicing pathway are also capable of impairing adenovirus replication.

To determine if these drugs showed similar specific activity on adenovirus mRNA splicing, we performed qRT-PCR on spliced and unspliced viral transcripts. To focus on the immediate effects of RNA splicing, A549 cells were infected with Ad5 before high- and low-doses of OTS and IsoG were subsequently added at 16 hpi and total RNA harvest at 24 hpi (**Figure 4I**). OTS behaved as a potent splicing inhibitor of both adenoviral early and late transcripts at the high dose (300 nM), but low dose (100 nM) treatment only had significant effects on late transcript splicing. IsoG had no significant or modest effects on splicing at the 30 μM dose, consistent with the above findings. At the higher 50 μM dose, however, IsoG strongly decreased the splicing of both adenoviral early E1A and late TPL splicing. Taken together, these data show that splicing inhibitors that impact alternative steps of the splicing cycle phenocopy many aspects of SF3B1-inhibition, including a preference for viral late transcription and protein production, decreased viral DNA replication, and strongly diminished infectious progeny production.

### Low-dose splicing inhibitors do not cause overt cytotoxicity

Splicing inhibitors have been previously reported to cause cell death in cancer cells, particularly those that also contain genetic mutation in spliceosomal components^26,27^. Cell health and viability is a critically important cofactor in productive viral infection. To address this potentially confounding variable, we performed 50% cytotoxic concentration assays (CC50) in A549 cells (**Supplementary Figure 2A**). A549 cells were subconfluently plated and allowed to grow in the presence of a wide dose range of all tested splicing inhibitors (PladB, H3B, OTS, IsoG). Global transcription inhibitor flavopiridol and cell-death inducing staurosporine were included as positive controls. After 48 hours of growth, total cell viability was measured by Alamar blue assay and normalized to vehicle DMSO treatment. In this assay, all tested compounds except IsoG had CC50s in the low nanomolar range. While potentially problematic, a metabolic readout such as Alamar blue cannot distinguish between cell death or slowed cell growth. Indeed, splicing inhibitors have been shown to alter cell cycle dynamics as well as predominantly causing cytotoxicity in a cycling-dependent manner^28^. To control for cell growth, we performed the same assay in immortalized primary Human Bronchial Epithelial Cells (HBEC3-KT). These cells were first grown to contact inhibition before being treated with the same concentration of drugs for 48 hours before percent viability was read out by Alamar blue (**Figure 5A**). In these non-dividing cells, a proper CC50 could no longer be calculated for three of the four splicing inhibitors (PladB, H3B, and IsoG), and OTS-964’s CC50 increased 10-fold to approximately 300 nM (**Supplementary Figure 2B**). HBECs were still sensitive to transcription inhibition or staurosporine-induced cell death. Considering viral infection also impairs cell cycle, it is likely that the previous effects on viral replication observed in the context of splicing inhibitors and after viral infection has been established are not explained by decreased cellular growth.

**Figure 5.**
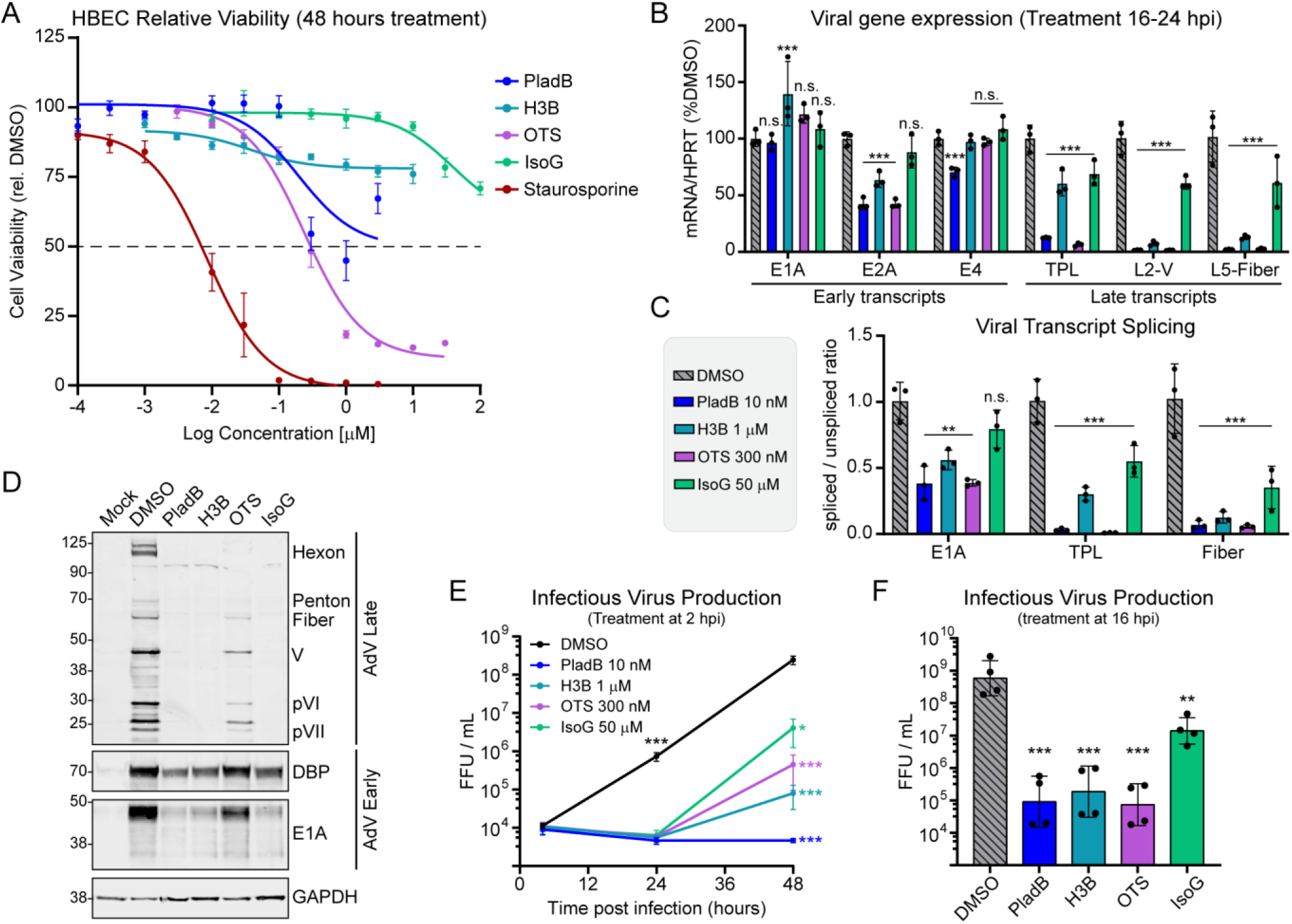
Splicing inhibition shows specific effects on viral replication in a primary cell model. **(A)** Human Bronchial Epithelial Cells (HBEC) were grown to contact inhibition in 96 well plates. Four independent biological replicates were treated with serial dilutions of PladB (blue), H3B (teal), OTS (purple), IsoG (green), or staurosporine (red) and incubated for 48 hours. Relative cell viability was measured by Alamar blue fluorescence assay and normalized to DMSO vehicle control at 100%. **(B)** Confluent HBECs were infected with Ad5 and subsequently treated with the listed doses of PladB, H3B, OTS, or IsoG at 16 hpi. Total RNA was collected at 24 hpi and qRT-PCR performed for viral transcripts normalized to HPRT and set to DMSO vehicle at 100%. **(C)** The unspliced transcript levels from B were calculated by qRT-PCR and the ratio of spliced to unspliced transcript is shown. For all qRT-PCR significance was analyzed by unpaired two-tailed *t*-test. **(D)** Immunoblot of uninfected confluent HBECs (Mock) or Ad5-infected confluent HBECs treated with vehicle control or 10 nM PladB, 1000 nM H3B, 100 nM OTS, or 50 μM IsoG at 2 hpi. Lysates were collected at 24 hpi and probed for Ad5 early and late proteins. GAPDH was used as a loading control. **(E)** Confluent HBECs were infected with fluorescent Ad5 (E3mNG) and treated with listed concentrations of splicing inhibitors at 2 hpi. Total cell plus supernatant was collected at 4, 24, or 48 hpi and released viruses used for virus progeny calculation via fluorescent forming unit (FFU). Data are presented as log-scale, points show the average of four biological replicates, and error bars show standard deviation. **(F)** Confluent HBECs were infected with fluorescent Ad5 with subsequent treatments of DMSO vehicle control or 10 nM PladB, 1000 nM H3B, 300 nM OTS, or 50 μM IsoG at 16 hpi. Total cell plus supernatant was collected at 48 hpi and released viruses used for virus progeny calculation via fluorescent forming unit (FFU). For all infectious virus statistical analyses significance was analyzed by lognormal *t*-test. Significance for all data was shown as P-value >0.5 (not-significant, n.s.), * P< 0.05, ** P< 0.01 and *** P< 0.001.

To investigate whether cell death or apoptosis played a larger role at the single cell level, we performed flow cytometry to detect apoptotic events after treatment with the highest used doses of splicing inhibitors (**Supplementary Figure 2C**). Cells were either treated with splicing inhibitors for 24 hours (**Supplementary Figure 2D**) or infected with Ad5 for two hours before subsequent 24 hour drug treatment (**Supplementary Figure 2E**). While both staurosporine and flavopiridol were able to induce apoptosis in infected or uninfected conditions, only 300 nM OTS-964 induced a statistically increased amount of apoptotic events, and only in the uninfected condition. Together, these data indicate that the effect of splicing inhibitors on viral infection is independent from cellular toxicity.

### Low dose splicing inhibition does not activate antiviral pathways

Previous reports have shown splicing inhibitors can cause cell death in cancer cells by inducing inflammation through sterile mimicry of viral infection. In these models, splicing inhibition causes retention of cellular transcript introns containing regions of repetitive double-stranded RNA that can activate RIG-I-like receptors and signal through downstream interferon pathways^29–31^. Such a finding could possibly explain the antiviral effects we see upon adenovirus infection through an indirect manner. To address this possibility, we performed qRT-PCR of uninfected cells or adenovirus infected cells that had been treated either with PladB, JAK/STAT kinase inhibitor Ruxolitinib, or both (**Supplementary Figure 3A**). Interferon-β treatment was included as a positive control upstream of interferon stimulated gene transcription (MX1, IFIT1, OAS1) as well as a control that Ruxolitinib was effective in dampening this ISG response. Not only did adenovirus infection induce very minimal or baseline ISG induction, as previously reported^32,33^, but PladB also did not induce interferon or ISG transcription. Ruxolitinib had no effect on either the infected or uninfected PladB-treated conditions. To assess whether any interferon effects downstream of JAK/STAT signaling outside of the tested ISGs could be affecting viral replication, we also performed a fluorescence focus forming assay of Ad5-infected A549 cells treated with PladB or concurrent PladB plus ruxolitinib (**Supplementary Figure 3B**). We once again saw a multiple-log decrease in infectious virus production in the presence of PladB, and the presence of ruxolitinib had no effect on this phenotype. Together, these data eliminate an indirect effect of antiviral induction as the mechanism for splicing inhibition’s effect on viral replication.

### Splicing inhibitors impair viral replication in non-transformed primary cells

We previously showed that all the tested splicing inhibitors have no gross cytotoxic effects in immortalized HBECs (**Figure 5A**). These same cells are also ideal models of adenovirus replication, and mimic the airway conditions and cells this serotype of adenovirus would encounter in a natural host. For all experiments HBEC cells were grown to contact inhibition before subsequent infection with high MOI 10 of Ad5 and subsequent treatment with effective doses of PladB, H3B, OTS, or IsoG ascertained in previous experiments. RNA was harvested 24 hpi and qRT-PCR was performed to assess total RNA (**Figure 5B**) and the relative splicing ratios of spliced and unspliced viral transcripts (**Figure 5C**). As in the transformed A549 cells, all tested drugs had minimal or non-significant effects on early transcripts E1A or E4, and less than 2-fold effects on E2A-DBP. In contrast, the total levels of late transcripts and splicing ratio of TPL and L5-Fiber were significantly and substantially downregulated. This preferential decrease in viral late proteins compared to early proteins was also seen by western immunoblot collected at 24 hpi (**Figure 5D**). Infectious virus production was measured by fluorescent focus forming assay over 24 and 48 hpi and all tested drugs induced multiple-log decreases in virus production (**Figure 5E**). Even when drug treatment was withheld to 16 hpi the amount of infectious virus production at 48 hpi was significantly reduced (**Figure 5F**). Compared to the 10-fold decrease seen with this same late treatment in transformed A549 cells, late treatment with IsoG decreased infectious virus production by almost 100-fold, and PladB, H3B, or OTS decreased infectious virus production by over 1000-fold. Together, these results indicate that the effect of splicing inhibitors on adenovirus is not cell-type or cancer cell-dependent.

### Splicing inhibition impairs Herpes Simplex Virus replication

Our initial results have demonstrated that inhibition of cellular splicing is incredibly potent on human adenoviruses that use cellular splicing machinery in the creation of nearly every viral transcript. We hypothesized that this sensitivity might extend to other human DNA viruses that also utilize the splicing machinery associated with RNA polymerase II. One such virus, Herpes Simplex virus 1 (HSV-1), can cause severe neurotropic and encephalitic infections^34^. Targeted antivirals such as acyclovir derivatives exist for these herpesvirus infections, but genetic mutations in the viral thymidine kinase or DNA polymerase can render these drugs ineffective^35^. Importantly, of the more than 100 characterized viral RNA transcripts, only five are spliced by cellular machinery. Some of these, such as the multifunctional ICP0 are important innate immune modulators but not necessary for viral infection^36^. Others, such as the terminase UL15, are essential for cleavage of the viral genome and packaging into capsids^37^. We set out to systematically test splicing inhibitors against this different DNA virus.

Normal human foreskin fibroblasts (HFF) were grown to contact inhibition and infected at high MOI 5 with HSV-1 strain Syn17. One hpi cells were treated with the previously determined effective doses of all four splicing inhibitors: 10 nM PladB, 1 μM H3B, 100 nM OTS, and 50 μM IsoG. The end of a single cycle of infection was collected at 16 hpi and immunoblotted for spliced ICP0 protein and unspliced control protein ICP8 (**Figure 6A**). All tested doses led to a decrease in the accumulation of ICP0 but had minimal effect on ICP8. Next we performed a flow cytometry experiment over a time course of infection with the recombinant Syn17 strain ND02 expressing YFP-tagged early gene ICP4 and RFP-tagged late gene VP26^38^. HFFs were infected at MOI 1.0, collected at 3, 6, 9, and 12 hpi, and analyzed for early and late gene expression (**Figure 6B**). All tested drugs decreased the accumulation of early YFP signal as early as 6 hpi and had a more pronounced decrease on late RFP accumulation as early as 9 hpi. In an orthogonal assay, the same virus was allowed to spread through a confluent culture of HFF cells by infecting at a low MOI of 0.01 per cell. Percent infection of the monolayer was determined by flow cytometry at 24, 48, and 72 hpi (**Figure 6C**). PladB and H3B significantly slowed down the spread of infection at 24 and 48 hpi. Both OTS and IsoG had very strong effects on this low MOI infection and essentially reduced viral replication to zero. Finally, production of infectious virus was quantified by plaque forming assay of the viruses produced during a 48-hour infection of HFF cells infected with a low MOI 0.01 spreading infection of HSV-1 (**Figure 6D**). PladB, H3B, and OTS led to significant decreases in viral production of approximately 10-20 fold, whereas IsoG led to a much stronger 3.5 log-fold decrease. This differential effect of PladB and IsoG was previously reported with a different strain of HSV-1 and in different cell types^11^. These results demonstrate that splicing inhibitors impair HSV-1 infection of primary cells, albeit with different kinetics and magnitudes of effect compared to adenovirus infection.

**Figure 6.**
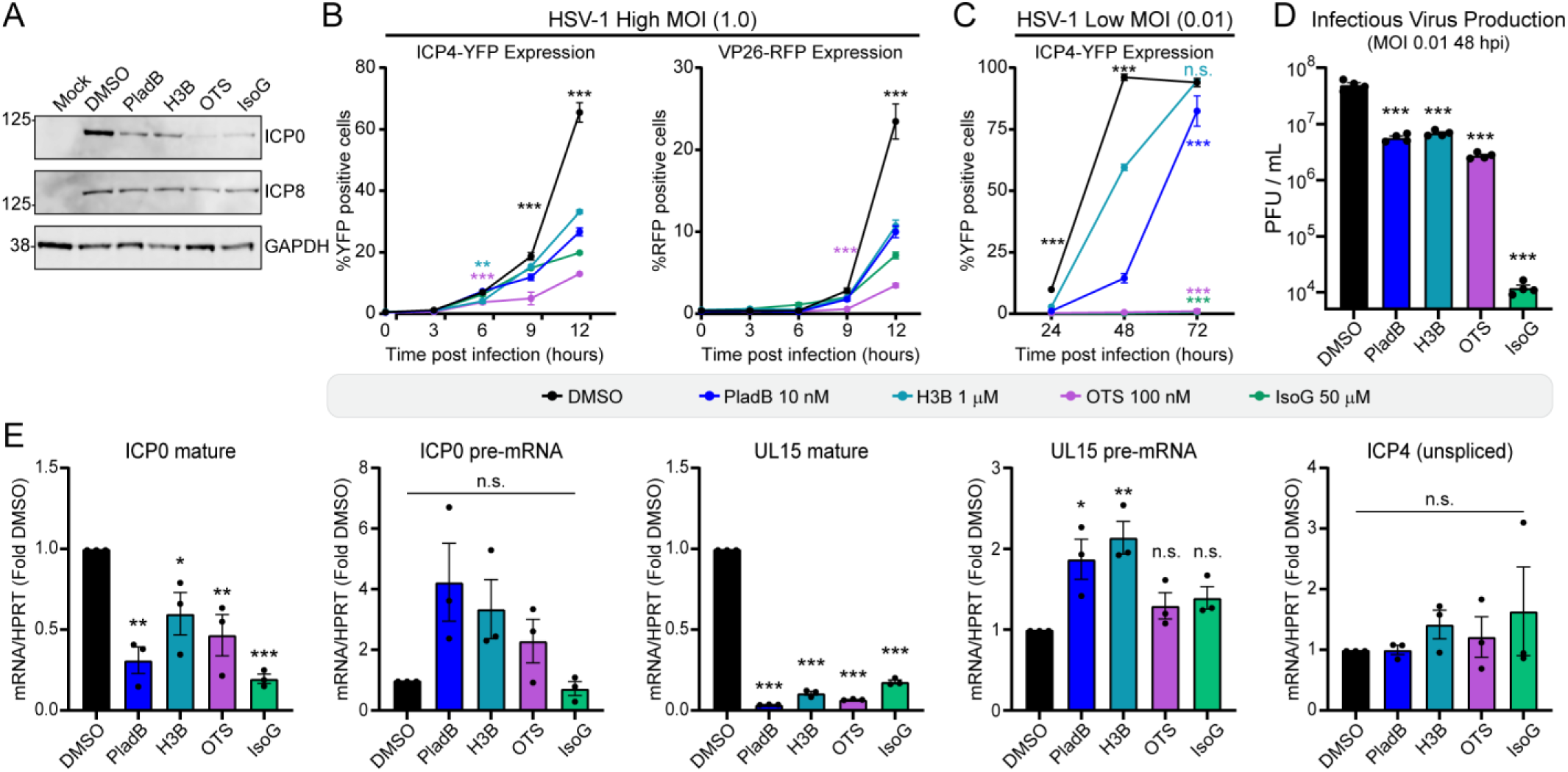
Splicing inhibition impairs viral RNA splicing and replication of Herpes Simplex Virus. **(A)** Immunoblot of uninfected human foreskin fibroblasts (Mock) or HSV-1-infected HFF cells treated with DMSO vehicle control or 10 nM PladB, 1000 nM H3B, 100 nM OTS, or 50 μM IsoG at 1 hpi. Lysates were collected at 16 hpi and probed for ICP0 (spliced product) or ICP8 (unspliced product). GAPDH was used as a loading control. **(B)** Confluent HFF cells were infected with fluorescent HSV-1 (ND02) at MOI 1.0 and treated with the listed concentrations of splicing inhibitors at 1 hpi. Cells were analyzed for percentage of cells positive for early YFP-ICP4 and late RFP-VP26 by flow cytometry at listed times. Data points represent the average of 3 biological replicates, and the error bars show standard deviation. **(C)** Confluent HFF cells were infected with fluorescent HSV-1 (ND02) at a low MOI of 0.01 and treated with the listed concentrations of splicing inhibitors at 1 hpi. Cells were analyzed for percentage of cells positive for YFP-ICP4 by flow cytometry at 24, 48, and 72 hpi. Data points represent the average of 3 biological replicates, and the error bars show standard deviation. **(D)** HFF cells were infected with HSV-1 at MOI 0.01 and subsequently treated with the listed concentration of splicing inhibitors at 1 hpi. Cells and supernatant were collected at 48 hpi and infectious progeny was measured by plaque assay upon Vero cells. Data are presented as log-scale, points show the average of four biological replicates, and error bars show standard deviation. **(E)** HFF cells were infected with HSV-1 at MOI 1.0 and subsequently treated with the listed concentration of splicing inhibitors one hour after infection. Total RNA was harvested at 8 hpi and qRT-PCR performed for spliced and unspliced ICP0 or UL15, as well as total ICP4. Transcripts were internally normalized to cellular HPRT as well as DMSO vehicle set to 1.0. Bars show the average of three independent biological replicates, and error bars show the standard deviation. For all qRT-PCR significance was analyzed by unpaired two-tailed *t*-test. Viral infectious titer was analyzed by lognormal ordinary one-way ANOVA. Flow cytometry was analyzed by two-way ANOVA and testing of comparisons per time point family. Significance for all data was shown as P-value >0.5 (not-significant, n.s.), * P< 0.05, ** P< 0.01 and *** P< 0.001.

We next assessed whether splicing inhibition was having on-target efficacy on HSV-1 transcript splicing. Confluent HFFs were infected with HSV-1 at MOI 1.0 and the same concentrations of all four drugs from above were added at one hour post infection. Total RNA was harvested at 8 hpi, and splice-specific qRT-PCR was performed for various HSV-1 transcripts (**Figure 6E**). Spliced transcripts included ICP0 and UL15 due to simplicity in distinguishing the relatively larger introns and excluding co-terminally spliced but separate genes^11^. For both transcripts, primers targeting the unspliced pre-mRNA isoform showed no decrease compared to vehicle control, and in fact had variable but sometimes non-significant upregulation. Splice-specific mature isoforms of both transcripts, however, substantially decreased in the presence of splicing inhibitor. Unspliced ICP4 transcript was included as a control, and its expression levels were not perturbed by drug treatment. Together, these data demonstrate that splicing inhibitors specifically impair the splicing of select HSV-1 transcripts that are spliced by cellular machinery, and that this inhibition leads to a significant decrease in viral replication.

### Replication of RNA viruses that undergo cell-mediated RNA splicing are impaired by splicing inhibitors

While most DNA viruses replicate in the nucleus using cellular machinery, RNA viruses rely on their own RNA-dependent RNA polymerases to perform processing of viral transcripts and genomes and primarily replicate in the cytoplasm. An important exception is Influenza virus, which replicates in the nucleus and has a complicated relationship with cellular RNA processing machinery^12,13^. Influenza virus is a negative sense RNA virus that enters cells pre-complexed with a multi-subunit RdRp that initiates transcription of viral mRNA from the individual eight genome segments using a “cap-snatched” primer derived from cellular transcripts^39^. Importantly, two of the Influenza segment mRNAs, M and NS, can undergo splicing mediated by cellular RNA binding factors and spliceosome components, including SF3B1, that occurs post-transcriptionally at nuclear speckles. The resulting transcripts M1 and M2, and NS1 and NEP, form critically important integral membrane proteins and innate immune antagonizing and replication proteins, respectively^40–43^. Post-transcriptional RNA splicing of these genes is utilized to regulate specific dosages of each gene transcript^44^. H1N1 and H3N2 strains of Influenza A Viruses (IAV) accumulate very different M1/M2 and NS1/NEP ratios^45,46^. While much work has targeted specific human splicing factors to understand the basic mechanisms of IAV post-transcriptional splicing, we hypothesized that untargeted and broad disruption of cellular splicing would likewise impair IAV replication.

A549 cells were infected with recombinant strains of pandemic 2008 California H1N1 or 2007 Brisbane H3N2 strains of IAV at an MOI of 0.1 PFU/cell. One hour post-infection, inoculation media was exchanged for viral growth media containing 10 nM PladB, 1 μM H3B, 100 nM OTS, or 50 μM IsoG. Forty-eight hours post infection supernatant was harvested, and the production of infectious virus was quantified by plaque assay (**Figure 7A-B**). For H1N1 virus both PladB and IsoG treatment reduced virus production by greater than 100-fold, while H3B still reduced virus production by an appreciable 14-fold. All drugs were slightly less efficacious for H3N2 infection, with both PladB and IsoG reporting approximately 20-fold reductions and H3B mediating a significant 5-fold reduction. OTS trended towards a modest effect for both viruses, but these results were not statistically significant. One explanation for these divergent results is that OTS impairs the kinase activity of CDK11 and likely impacts splicing co-transcriptional with RNA polymerase II, which is a very different mechanism from the post-transcriptional splicing employed by Influenza^21,22^. These results show that splicing inhibitors that can target RNA processing distinct from RNA polymerase II can functionally impair virus production from two clinically relevant strains of IAV.

**Figure 7.**
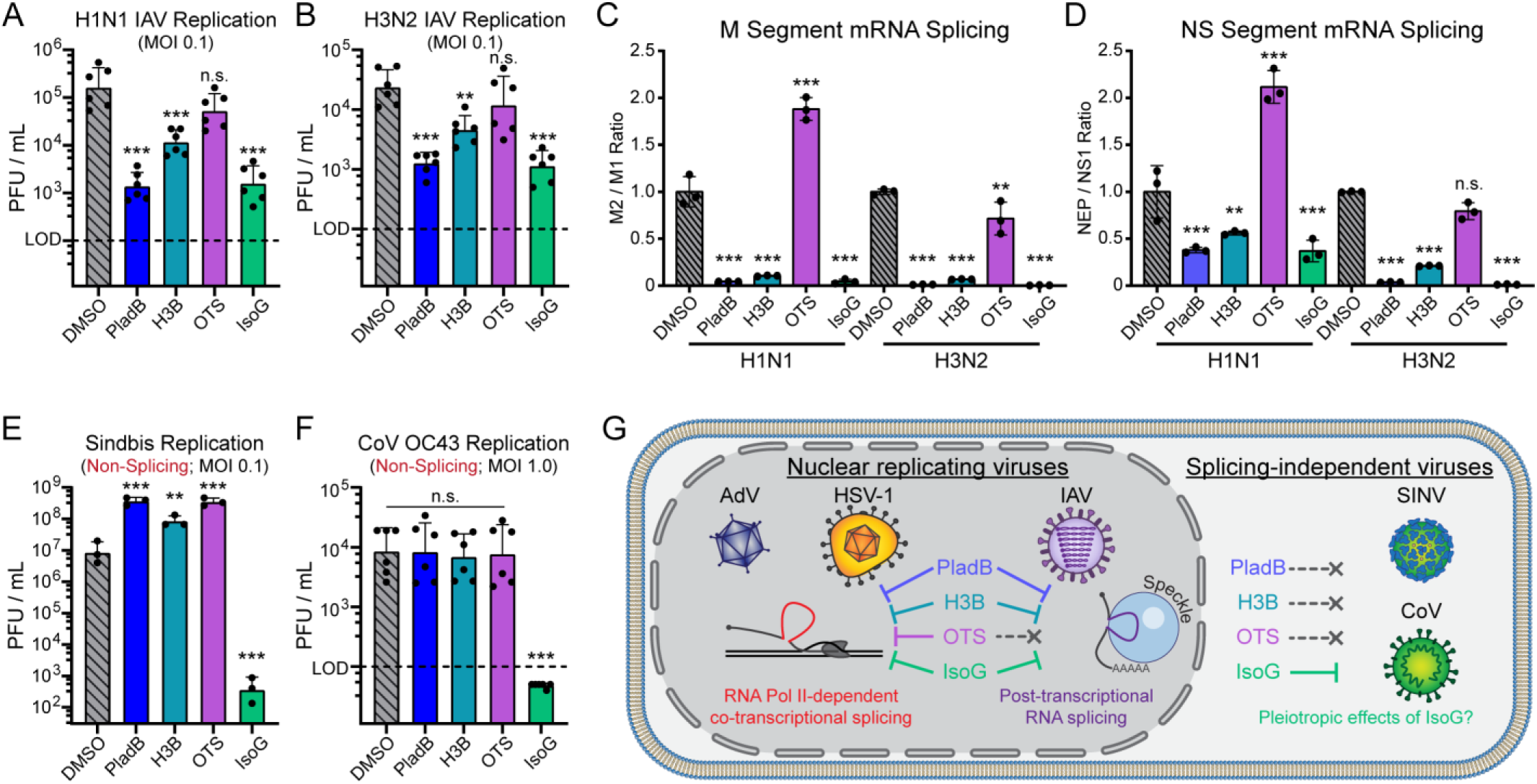
Splicing inhibition is effective against diverse RNA viruses that undergo cellular-dependent RNA splicing. **(A)** A549 cells were infected with recombinant 2008 pandemic H1N1 IAV at MOI 0.1 and subsequently treated with 10 nM PladB (blue), 1000 nM H3B (teal), 100 nM OTS (purple), or 50 μM IsoG (green) at 1 hpi in media containing BSA and TPCK trypsin. At 48 hpi cell supernatant was collected and titered on MDCK cells by plaque assay. Data are presented as log-scale, points show two independent experiments with six experimental replicates, and error bars show standard deviation. **(B)** Same as in A, but A549 cells were infected with recombinant 2007 Brisbane H3N2 IAV. **(C)** A549 cells were infected with H1N1 or H3N2 IAV at MOI 1.0 and treated with the same dosages of splicing inhibitors at 1 hpi. Total RNA was harvested at 24 hpi and reverse-transcribed using oligo-dT primers. qRT-PCR was performed using splice-specific primers for M1 and M2 transcripts, and the M2/M1 ratio was reported for three independent biological replicates. Bars show average of all three replicates, and error bars show standard deviation. **(D)** Same as in C, but total mRNA was analyzed by qRT-PCR and splice ratios constructed for NEP/NS1 ratio of transcripts from the NS segment. **(E)** A549 cells were infected with Sindbis virus at an MOI of 0.1 and subsequently treated with 10 nM PladB (blue), 100 nM H3B (teal), 100 nM OTS (purple), or 50 μM IsoG (green) at 1 hpi. At 24 hpi supernatant was harvested and titered on Vero cells by plaque assay. Data are presented as log-scale, points show the average of three biological replicates, and error bars show standard deviation. **(F)** A549 cells were infected with OC43 coronavirus at an MOI of 1.0 and subsequently treated with 10 nM PladB (blue), 1000 nM H3B (teal), 100 nM OTS (purple), or 50 μM IsoG (green) at 1 hpi. At 24 hpi supernatant was harvested and titered on vero cells by plaque assay. Data are presented as log-scale, points show two independent experiments with six experimental replicates, and error bars show standard deviation. **(G)** Unifying model explaining the divergent effects of splicing inhibitors on both splicing-dependent and non-splicing viruses. All qRT-PCR significance was analyzed by unpaired two-tailed *t*-test. Viral infectious titer was analyzed by lognormal ordinary one-way ANOVA. Significance for all data was shown as P-value >0.5 (not-significant, n.s.), * P< 0.05, ** P< 0.01 and *** P< 0.001.

We next assessed whether splicing inhibition was having on-target efficacy on IAV post-transcriptional splicing. A549 cells were infected with H1N1 or H3N2 IAV at MOI 1.0 and treated with the same concentrations of all four drugs at one hour post infection. Total RNA was harvested at 24 hpi, reverse-transcribed with oligo-dT primers to specifically enrich IAV mRNA transcripts, and splice-specific qRT-PCR was performed for M1, M2, NS1, and NEP transcripts (**Figure 7C-D**). Plotting the splice ratio of M2/M1 or NEP/NS1 revealed that PladB, H3B, and IsoG all led to the specific decrease of the spliced RNA isoform and therefore a decrease in total splicing ratio. H1N1 splicing was more strongly affected compared to H3N2, mirroring the above data on virus replication, and is probably related to the previously reported differences in splice site strength and relative ratios between the two strains^45,46^. OTS did not decrease the splicing ratio for either transcript pair in either strain. These data demonstrate that inhibitors targeting machinery shared between co- and post-transcriptional splicing processes can impair the splicing processes employed by evolutionary distinct influenza viruses leading to a decrease in viral replication.

We previously demonstrated that interferon transcripts do not appear to be induced by splicing inhibition (**Supplementary Figure 3**); however, potential confounding effects due to alterations in cell metabolism or other processes can still affect downstream viral production. To address these concerns, we infected A549 cells with two distinct cytoplasmic replicating RNA viruses that do not undergo any form of cell-based RNA splicing. We hypothesize that if splicing inhibition alters a general pathway necessary for viral infection, we would see a decrease in the infectious virus production of these non-splicing viruses.

We first utilized Sindbis virus (SINV), an arthropod-borne alphavirus that was recently shown to be inhibited by SF3B1 directly in a non-splicing manner^47^. We recapitulated these findings with PladB and IsoG, as well as expanded the scope to include clinical-grade SF3B1 inhibitor H3B and SF3B1 phosphorylation inhibitor OTS (**Figure 7E**). Our results fully phenocopy and expand on the results of Kamel et al, showing approximately 10-fold increased Sindbis replication upon treatment with any of the SF3B1-targeting inhibitors, as well as substantial impairment upon 50 μM IsoG treatment. Finally, we investigated infection with OC43, a coronavirus that circulates seasonally and causes a common cold phenotype^48^. Coronaviruses employ discontinuous genome transcription to produce subgenomic mRNA that structurally look like spliced RNAs, however the viral mechanism for producing these transcripts is functionally distinct from, and does not rely on, cellular spliceosomes^49^. A549 cells were infected with OC43 at an MOI of 1.0 PFU/cell, treated with splicing inhibitors one hpi, and supernatants were assayed at 24 hpi by plaque assay. While PladB, H3B, and OTS had no significant effect on OC43 replication, 50 μM IsoG treatment was highly antiviral and reduced infectious titer below the limit of detection. Taken together, these data imply that whatever off-target effects SF3B1-inhibition has in cells outside of splicing inhibition, these are not sufficient to impair the ability of two distant RNA viruses to complete their full cycle of infection. The results with Isoginkgetin are potentially more puzzling. Our data shows high doses (> 30 μM) of IsoG are capable of inhibiting cellular, adenoviral, HSV, and influenza splicing. However, recent publications have shown that IsoG is a poor splicing inhibitor^50^. In addition, IsoG has been reported to have many additional effects, including proteosome inhibition and decreased protein synthesis^51,52^. We believe it is likely that IsoG might have an anti-splicing effect against splicing-dependent viruses, as well as other potent off-target effects that affect viral replication more broadly.

A unifying model summarizing these relationships across viral families is presented in **Figure 7G**. We have shown that four mechanistically distinct splicing inhibitors demonstrate antiviral activity across three evolutionarily divergent viral families. Viral sensitivity tracks with each virus’s reliance on host-mediated RNA processing, while cytoplasmic RNA viruses lacking spliceosome dependence remain unaffected, supporting a direct, on-target mechanism. Together, these findings establish host spliceosome dependence as a conserved and pharmacologically tractable vulnerability of nuclear-replicating viruses.

## Discussion

Our results establish splicing inhibition as a viable host-targeted therapeutic strategy against nuclear-replicating viruses. In adenovirus, splicing inhibitors produced a stage-specific effect, with late viral transcripts and proteins consistently more sensitive than their early counterparts. This effect was recapitulated by genetic depletion of SF3B1 and by mechanistically distinct compounds (OTS-964, Isoginkgetin), indicating that the antiviral phenotype reflects a general dependence on cellular splicing rather than inhibition of any single step in the splicing cycle. This vulnerability extends to several viruses that rely on host-mediated RNA processing, underscoring the breadth and therapeutic potential of this mechanism. For HSV-1, splicing inhibitors impaired infection from the immediate-early stage onward, coincident with the onset of expression of the small subset of spliced viral transcripts. A related effect of Pladienolide B and Isoginkgetin was reported for HSV-1 canonical, but not circular, RNA splicing^11^. For IAV, splicing inhibitors strongly suppressed generation of the spliced M2 and NEP isoforms across two distinct strains, with corresponding defects in viral replication. A prior study reported that a different SF3B1-targeting compound, herboxidiene, suppresses M2 splicing^46^. Our results extend this observation to three additional, mechanistically distinct inhibitors, and demonstrate a parallel effect on NS1/NEP splicing. Importantly, we show that the CDK11 inhibitor OTS-964 is largely ineffective against influenza, likely reflecting differences between Pol II-mediated co-transcriptional splicing and influenza post-transcriptional splicing^14^. Together, these findings indicate that the splicing-dependent vulnerability identified in adenovirus extends across evolutionarily diverse viral families, with the degree of antiviral sensitivity tracking the extent to which each virus relies on RNA splicing.

In contrast, splicing-independent cytoplasmic viruses were not impaired by splicing inhibition. In the case of Sindbis virus, SF3B1 and CDK11 inhibitors modestly enhanced replication, consistent with a previously described moonlighting role for splicing factors in restricting this specific virus^47^. OC43 coronavirus replication was largely unaffected by these compounds, which contrasts with an earlier drug-screening study identifying Pladienolide B as a candidate anti-coronaviral agent^53^. This discrepancy may reflect differences in drug concentration or cell-intrinsic effects. Notably, the antiviral effects described throughout our study were observed at doses that preserved host cell viability, arguing against general cytotoxicity as a confounding explanation for impaired viral replication. Previous reports have observed splicing inhibition triggering antiviral interferon induction through cellular dsRNA associated with intron retention^29–31^; however, we observed minimal interferon pathway activation under our cellular and dosage conditions. Together, these important controls indicate that the antiviral activity of splicing inhibitors reflects a direct, on-target consequence of disrupted spliceosome function rather than indirect effects on host cell health or innate immune signaling.

Our data also highlights a potentially confounding distinction between splicing-dependent and splicing-independent effects of isoginkgetin. Unlike pladienolide B, H3B-8800, and OTS-964, whose antiviral activity correlated closely with their effects on splicing across all tested viruses, isoginkgetin retained activity against splicing-independent viruses at the high concentrations needed to achieve efficacy on cellular RNA splicing. This could be attributed to isoginkgetin’s off-target effects on the proteasome, protein synthesis, or general transcription being greater than the effect of splicing inhibition for these viruses^50–52^. We therefore interpret the antiviral activity of isoginkgetin with caution, and consider the concordant results obtained with the three SF3B1-and CDK11-targeting compounds to provide the strongest evidence for splicing inhibition as a direct antiviral mechanism.

While potent direct-acting antivirals exist for both herpesviruses and influenza, these drug classes remain vulnerable to resistance-conferring viral escape mutations^35,54^. Host-targeted antivirals offer an attractive alternative because host targets cannot be readily mutated by the virus^55,56^. However, this strategy has historically been limited by concerns over toxicity when targeting essential cellular machinery. Adenovirus remains a serious cause of morbidity in immunocompromised and pediatric populations, yet no approved antiviral or vaccine exists^57–59^. Several groups have explored repurposing existing compounds against adenovirus^60–62^, most notably the CDK9 inhibitor flavopiridol, which broadly suppresses RNA Polymerase II-dependent transcription and shows a similar therapeutic window against adenovirus in cell culture^63,64^. However, despite extensive clinical development across multiple cancer types, flavopiridol has consistently been limited by severe, dose-limiting toxicity *in vivo*^65^. Splicing inhibitors may offer a more tractable path forward compared to inhibition of transcription itself, as these inhibitors appear to spare the bulk transcriptional output required for normal cell function while still disrupting the alternative processing events on which complex viral transcripts disproportionately depend. More broadly, the growing clinical pipeline of splicing modulators developed for spliceosome-mutant malignancies provides a ready-made toolkit of compounds with established pharmacokinetics and safety data in humans^4^. This idea of repurposing anti-cancer compounds for antiviral effect is not without precedent: Azidothymidine (AZT) was first developed as a cancer therapeutic before its repurposing as the first antiretroviral approved for HIV^66^. Our findings suggest that this pipeline merits evaluation not only for its original oncology indications, but also as a source of candidate broad-spectrum antivirals against nuclear-replicating viruses.

## Acknowledgments

We thank M. Weitzman, B. Tian, P. Lieberman, and all members of the Price lab for careful review of our manuscript. This research was supported by NIH NIAID grants R00-AI159049 (A.M.P.), R01-AI186810 (D.T.C.), R01-AI140442 (S.R.W.), R01-AI169537 (S.R.W.), NIAID HIV Pathogenesis training grant T32-AI007632 (R.W.A.), and NIH NCI national research service award training grant T32-CA009171 (A.Y.). Additional support came from the Wistar Science Discovery Fund and Goldblum Family Healthcare Fund, as well as the W.W. Smith Charitable Trust grant #C2506 (A.M.P.). Wistar Shared Resources are supported by NIH grant P30-CA010815. We thank the Wistar Flow Cytometry Shared Resource for help with flow cytometry analysis.

## Materials and Methods

### Cell Culture

A549 (CCL-185), HFF-1 (SCRC-1041), U2OS (HTB-96), and VeroE6 cells (CRL-1586) were obtained from ATCC. AD-293 (240085) were obtained from Agilent. Vero cells were a generous gift from M. Weitzman. MDCK cells were a generous gift from S. Hensley. All of the above cells were cultured in Dulbecco’s modified Eagle medium (DMEM; Corning, 10-013-CV) supplemented with 10% vol/vol fetal bovine serum (Cytiva, SH30910.03HI) and 1% vol/vol pen-strep (Corning, 30-002-CI) at 37°C and 5% CO_2_, except HFF which were cultured in 15% vol/vol fetal bovine serum. Immortalized Human Bronchial Epithelial Cells (HBEC3-KT, CRL-4051) were obtained from ATCC and grown in Airway Epithelial Cell Basal Medium (PCS-300-030) supplemented with HLL supplement, L-Glutamine, Extract P, Airway Epithelial Cells Supplement (PCS-300-040) and 0.1% vol/vol pen-strep (Corning, 30-003-Cl) as described by the manufacturer. All cell lines tested negatively for mycoplasma contamination using the MycoAlert Mycoplasma Detection Kit (Lonza, LT07-418).

### Viral Infections

Wildtype adenovirus serotype 5 (Ad5) was created using AdenoBuilder transfection system as previously described^23,67^. Fluorescent adenovirus reporters E3mNG and E3gL5s were described previously^23^. All adenoviruses were expanded using AD-293 cells and purified using two sequential rounds of ultracentrifugation in CsCl gradients and stored in 40% vol/vol glycerol at - 30°C (short term) or -80°C (long term). Infections with Adenovirus were performed at a multiplicity of infection (MOI) of 10 FFU/cell. Cells were infected at 80-90% confluent monolayers by incubation with diluted virus in a minimal volume of low serum media (2%) for 2 hours.

Dual color HSV-1 reporter virus ND02 (YFP-ICP4 / RFP-VP26) was obtained from N. Drayman^38^. Virus was expanded in Vero cells and titered via plaque assay in U2OS cells. Spreading infections were performed at a multiplicity of infection (MOI) of 0.01 PFU/cell, while other infections were performed at an MOI of 1-5 PFU/cell. Infections of 100% confluent monolayers of HFF cells were performed by incubation with diluted virus in a minimal volume of low serum media (2%) for 1 hour.

P0 recombinant WT H1N1 (A/California/04/2009 H1N1 (pH1N1)) was provided by Dr. Luis Martinez-Sobrido^68^. Clinical isolate of seasonal H3N2 (A/Brisbane/10/2007) was provided by S. Hensley to Susan Weiss. Both influenza strains were passaged once on MDCK cells at MOI = 0.001 to produce P1 virus stock which was used for all infections. A549 cells were washed 1X with PBS followed by infection with H1N1 at MOI = 0.1 at 37°C for 1 hour with rocking every 15 minutes. After 1 hour for viral adsorption, cells were washed 1X with PBS and supplemented with RPMI containing 0.2% BSA, 1μg/mL TPCK-treated trypsin, and inhibitor or DMSO at indicated concentration. Supernatant was collected at indicated time post infection followed by quantification of viral titer via plaque assay on MDCK cells.

GFP-expressing Sindbis virus was a generous gift from M. Heise to Susan R Weiss^69^. The virus was expanded in Vero cells in 2% DMEM media and stored at -80 °C (long term storage); thawed aliquots were stored for up to 1 week at 4°C. Viral stock titer was determined by plaque assay on Vero cells using Avicel 0.6% overlay. Sindbis infections were performed at a MOI of 0.1 PFU/cell. Cells were infected at 80-90% confluent monolayers by incubation with diluted virus in a minimal volume of low serum (2%) for 1 hour.

OC43 coronavirus was obtained from ATCC (VR-1558) and expanded on VeroE6 cells at 33°C. Infections were performed on A549 cells washed 1x with PBS then infected at MOI 1 PFU/cell in serum-free RPMI media at 33°C for 1 hour. After viral adsorption, cells were washed 1X with PBS followed by addition of RPMI containing 2% FBS + drug at respective concentration. Supernatant collected at indicated times post infection, followed by quantification of viral titer via plaque assay on VeroE6 cells with 5 day incubation at 33°C followed by fixation in 4% PFA and visualization of plaques with crystal violet.

### Infectious titer calculations

Viral titers for adenovirus infections were determined using a fluorescence-based assay employing an AdV reporter construct containing mNeonGreen in the viral E3 transcriptional unit. The detailed procedure is described as previously^23^. Briefly, total infected cells plus supernatant were harvested at a given time point and frozen at -80 °C. Lysate was freeze/thawed thrice in dry ice/methanol baths with vortexing between thaws. Cell debris was pelleted by centrifugation at 10,000 RCF and cleared lysates were transferred to new tubes. Infectious virus was serially diluted in 2%DMEM media into 96-well plates containing a monolayer of A549 cells. At 48 hpi plates were imaged on a LiCor Odyssey M using the 488 laser and fluorescent puncta were counted using the Find Maxima tool of ImageJ.

All other viruses were quantified by plaque assay. HSV-1 infected cells were collected at the desired timepoint by removing the supernatant (cell-free virus) and cell-associated virus separately. Cells with cell-associated virus were freeze/thawed thrice and pre-cleared using the method above before being recombined with the cell free supernatant and frozen at -80 °C. Monolayers of U2OS cells were infected with serial dilutions of HSV-1 in 2%DMEM for two hours before washing with PBS and overlay with methocel media (1:1 mixture of 2% methylcellulose solution in water and 2X DMEM). At 3 days post infection overlays were aspirated, cells were washed with PBS, and then fixed with a solution of 1% Crystal Violet (Millipore Sigma, C6158-50G) and 50% methanol.

### Western Blotting

For western blotting, protein samples were lysed in RIPA buffer (1% IGEPAL CA-630, 0.1% SDS, 150mM NaCl, 50 mM Tris (pH 7.4)) supplemented with 1X Halt Protease Inhibitor (Fisher Scientific, P178430) and quantified by BCA assay (Thermo Scientific, 23225). For gel electrophoresis, 10-15 μg (for AdV) or 30-40 μg (for HSV-1) of proteins were mixed with 1:4 NuPage LDS, supplemented with 1 mM DTT. Lysed samples were boiled at 95°C for 15 minutes, cooled, then vortexed and spun down before loading. Proteins were separated in precast NuPAGE polyacrylamide gels, or self-cast using the Invitrogen SureCast Gel Handcast System, and transferred onto nitrocellulose membranes (Bio-Rad, 1620115). Membranes were blocked with 5% non-fat milk (Lab Scientific, M0841) plus 0.05% Sodium Azide (Millipore Sigma, S2002-500G) and then rocked in primary antibody overnight at 4°C. The following primary antibodies were utilized: Anti-Ad5 Capsid at 1:5,000 (Abcam, ab6982), E1A at 1:500 (BD Biosciences, 554155), DBP at 1:500 (hybridoma supernatant, mouse clone B6-8, original from A. Levine, kind gift of M. Weitzman), ICP0 at 1:1000 (Santa Cruz Biotechnology, sc-53070), ICP8 at 1:1000 (Abcam, ab20194), and GAPDH at 1:2,000 (LiCor, 926-42216).

Membranes were washed three times with 1× TBST (Tris-Buffered Saline with Tween 20; 10 mM TrisHCl, 0.01% Tween 20, 150 mM NaCl in MilliQ Water), then anti-rabbit (LiCor, 926-68171), or anti-mouse (LiCor, 827-08364) fluorescent secondary antibodies at 1:10,000 in TBST were applied to the membranes for a period of two hours. Alternatively, anti-mouse (Genetex, GTX213111-01) HRP antibody at 1:10,000 in TBST was applied to the membranes for a period of two hours. Membranes were washed three more times with 1× TBST and scanned on a LiCor Odyssey M using either fluorescent or chemiluminescent settings.

### RNA transfection and knockdown

The following siRNA pools were obtained from Dharmacon: siGENOME Human SF3B1 (23451) siRNA (M-020061-02-0005), and siGENOME Non-Targeting siRNA Pool #1 (D-001206-13-20). All siRNA transfections were performed using the Lipofectamine RNAiMAX (Invitrogen) protocol. Transfections were performed on A549 cells for 24 hours and cells were subsequently infected with wildtype Ad5 or mock for 24 hours as described above.

### Fluorescent imaging and CellCyte live cell imaging

A549 cells were infected with the fluorescent AdV reporter E3gL5s at a MOI of 10 FFU/cell. This construct contains mNeonGreen in the viral E3 transcriptional unit and a cleavable P2A-mScarlet3 into the L5-Fiber late region. The fluorescent reporter was imaged on an EVOS imager (ThermoFisher) at 24 hpi. The same reporter construct was also used to perform CellCyte live cell imaging. A549 cells were seeded in a single monolayer at 80%–90% confluence in black 96-well plates (Greiner, 655090) and infected with E3gL5s at an MOI of 10 FFU/cell for 2 hours before adding drug treatments and beginning live cell imaging at 6 hpi. Using a 4× objective and four images per well, green and red fluorescence were captured every hour for 48 hours and analyzed using CELLCYTE X software to threshold and determine the total number of fluorescent objects per well.

### RNA and DNA isolation

Total RNA was isolated from cells by RNeasy Mini Kit (Qiagen, 74106), following manufacturer protocols. RNA was treated with RNase-free DNase I (Qiagen, 79256) on-column. 500 ng or 1 µg of input RNA were reverse transcribed using the High-Capacity cDNA Reverse Transcription Kit (ThermoFisher, 4368814). For Influenza A virus, reverse transcription was performed combining the High-Capacity cDNA Reverse Transcription Kit with Oligo(dT)_16_ primers (Invitrogen, 100023441). Total DNA was crudely isolated by resuspending total cell pellets in a 5% Chelex solution (Sigma Aldrich, C7901) in TE Buffer before boiling samples at 95°C for 15 minutes and subsequent incubation at RT for 5 minutes. Samples were centrifuged at 12000 RCF for 90 seconds at RT and clean supernatant was collected and DNA was quantified by nanodrop.

### Quantitative PCR

Quantitative Reverse Transcription PCR (qRT-PCR) was performed combining 10 ng of cDNA with 10 μM forward and reverse primers (**Supplementary Table 1**), and 2xPowerUp SYBR Green Master Mix (Applied Biosystems, A25742), in a 12 μL reaction. DNA qPCR was performed by performed combining 20-40 ng of DNA, together with 10 μM primer pairs (either Tubulin or DBP), and 2xPowerUp SYBR Green Master Mix in a 12 μL reaction. QuantStudio 6 Flex System. All values were normalized by ΔΔCt method normalizing to HPRT as an internal control for RNA or Tubulin as a control for DNA.

### Semi-quantitative RT-PCR

PCR amplification was performed on 10 ng of cDNA (or 10 ng of genomic DNA control) in 20 µl of reaction mixture including 1x Phusion HF Buffer, 200 μM dNTPs, 1 μM of Forward and Reverse primers and 1.0 units/50 µl of Phusion DNA polymerase (NEB, M0530S). PCR conditions for DNAJB1, BRD2, and GAPDH primers were: 94°C for 5 minutes; 35 cycles of 94°C for 30 sec, 60°C for 30 sec, 72°C for 1 min; and final extension at 72°C for 5 minutes. PCR products were stained with GelRed Prestain Loading (Biotium, 41010) and ran on a 2% agarose gel at 120 V for 45 minutes. Molecular weights of the amplification products were quantified using 1 Kb Plus DNA Ladder (Invitrogen, 10787018).

### Alamar Blue cell viability assay

Cell viability assay was performed using the AlamarBlue^TM^ Cell Viability Reagent (Invitrogen, A50101), seeding conditions varied among cell lines. A549 cells were seeded counting 5000 cells/well in a 96-well plate and treated 1 day after seeding. HBECs were seeded at 10000 cells/well and let reach total confluence before proceeding with treatments. 1 or 2 days after seeding, culture media was aspirated and fresh media was added containing serial dilutions of drug treatments. After 48 hours, 1/10th of cell viability reagent was added to each well; plates were incubated for 1 hour at 37°C before measuring fluorescence (ex. 560 nm, em. 590 nm) or absorbance (570 nm) on a BioTek Synergy Neo2 multi-mode plate reader.

### Apoptosis assay and flow cytometry

100,000 A549 cells were plated per well in 12-well plates 24 hours prior to treatment, and then fresh media was replaced containing drug doses for an additional 24 hours. After treatment, cells were trypsinized and half volume of the resuspended cells was transferred in flow cytometry tubes to perform cell death assay. Cells were centrifuged at 300 RCF for 5 minutes and washed with 1x Annexin Binding Buffer twice, then were resuspended in 100 μL of 1x Annexin Binding Buffer. In each tube were added 5 μL of Alexa Fluor 488 conjugated Annexin V (Invitrogen, A13201) and 1 μL of Propidium Iodide (Invitrogen, V13241B) and incubated RT for 20 minutes. Samples were processed on a FACSymphony A3 Cell Analyzer (BD Biosciences) and analyzed using FlowJo Software (BD Biosciences).

### Drugs and small molecule inhibitors

All drugs were resuspended in Dimethyl Sulfoxide (DMSO) and frozen at -20°C in multiple aliquots to allow for low numbers of freeze/thaw cycles. Pladienolide B was purchased from Santa Cruz Biotechnology (sc-391691). H3B-8800 was purchased from Cayman Chemical (42106) and Medinoah (90-7523). OTS-964 was purchased from Cayman Chemical (17052). Isoginkgetin was purchased from Millipore Sigma (41615410MG). Flavipiridol (S1230) and Staurosporine (S1421) were purchased from Selleck Chemicals. Ruxolitinib was purchased by Cayman Chemical (11609). Human Interferon-β was purchased from GenScript (Z03109) and resuspended in BSA 1% PBS, then stored in -20°C.

---

**Supplementary Figure 1.**
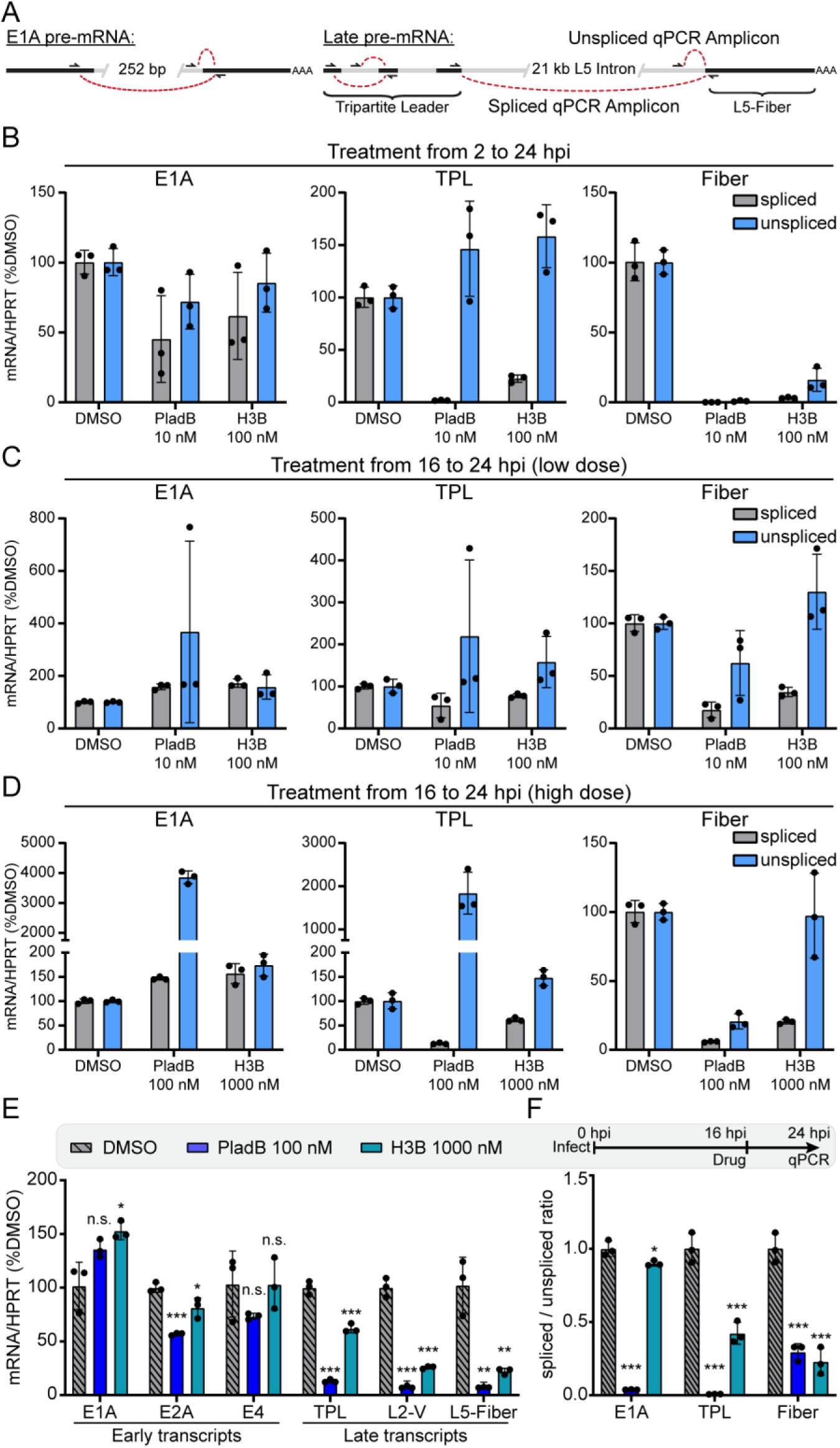
Spliced-transcript aware PCR reveals specific effect of both low- and high-dose splicing inhibition. **(A)** Model showing representative transcripts and primer design to identify spliced and unspliced adenoviral early and late transcripts by qRT-PCR. **(B)** Individual spliced and unspliced E1A, TPL, and Fiber transcript results used to calculate the splicing ratio in Figure 2B. **(C)** Individual spliced and unspliced E1A, TPL, and Fiber transcript results used to calculate the splicing ratio in Figure 2D. **(D)** A549 cells were infected with Ad5 and treated with higher concentrations of PladB (100 nM, blue) or H3B (1000 nM, teal) at 16 hpi. Total RNA was harvested at 24 hpi and individual spliced and unspliced E1A, TPL, and Fiber transcripts are shown. **(E)** Total RNA from D was analyzed by qRT-PCR for listed viral early and late transcripts and normalized to both cellular HPRT1 transcripts and DMSO-treated vehicle controls. Data points are shown for three biological replicates, bars depict mean, and error bars show standard deviation. **(F)** Data from D rearranged to show the splicing ratio of E1A, TPL, and Fiber. Significance was analyzed by unpaired two-tailed *t*-test, and displayed as P-value >0.5 (not-significant, n.s.), * P< 0.05, ** P< 0.01 and *** P< 0.001.

**Supplementary Figure 2.**
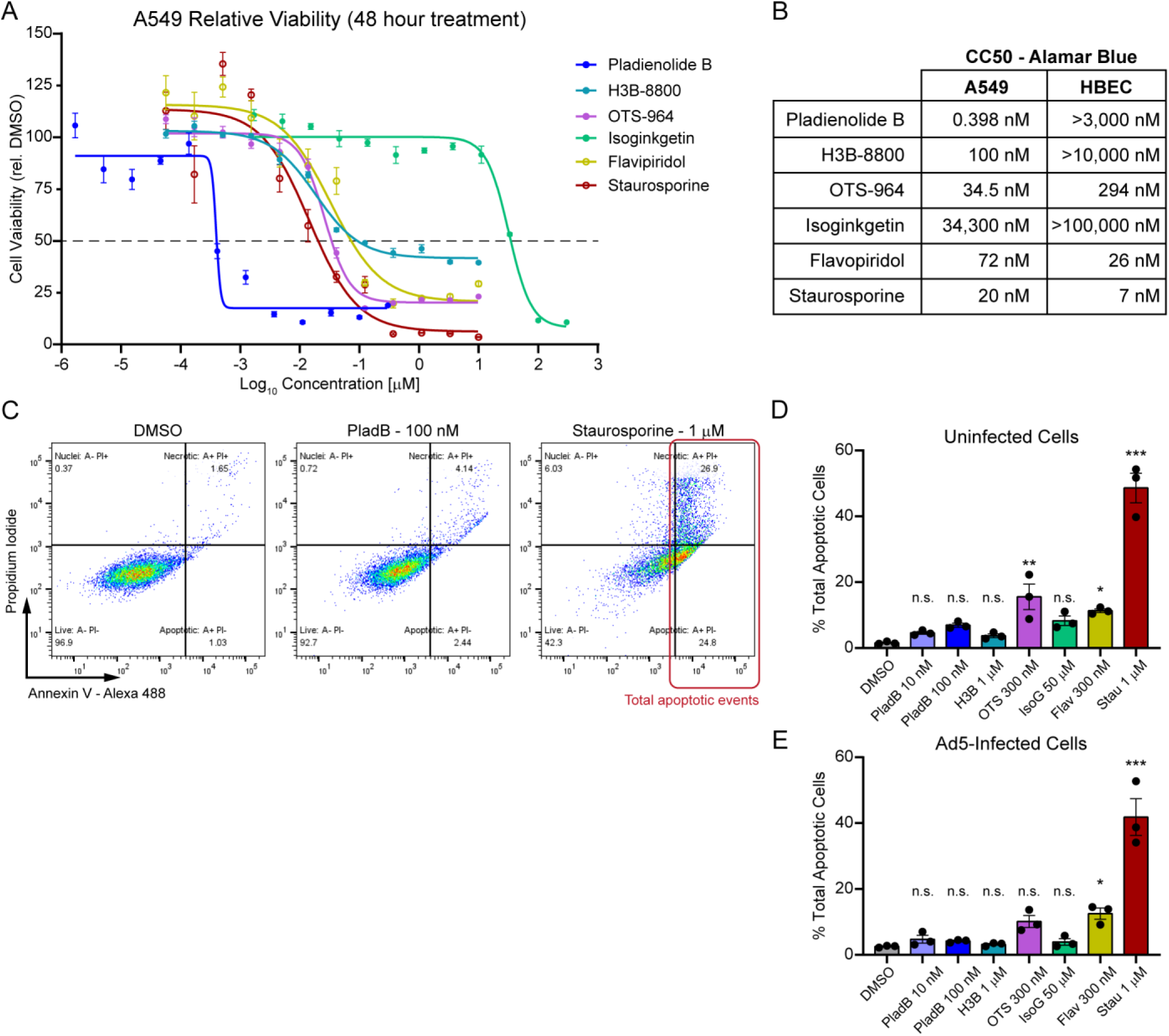
Low-dose splicing inhibitors do not cause overt cytotoxicity. **(A)** A549 cells were seeded in 96 well plates and treated with serial dilutions of PladB (blue), H3B (teal), OTS (purple), IsoG (green), flavopiridol (yellow), or staurosporine (red) and incubated for 48 hours. Relative cell viability was measured by Alamar blue fluorescence assay and normalized to DMSO vehicle control at 100% and media only at 0%. Points represent four independent biological replicates, and error bars denote standard deviation. Curve was fitted using a four parameter log(inhibitor) vs response formula. **(B)** Cytotoxic Concentration at 50% (CC50) values constructed using Alamar blue viability data from the 50% intercept of the calculated nonlinear fit curve for each drug for both A549 cells and the confluent HBECs assayed in Figure 5A. **(C)** Representative flow cytometry plots for Annexin V and propidium iodide-based cell apoptosis assay performed on uninfected A549 cells treated with the listed concentration of drugs for 24 hours. **(D)** Total apoptotic events (Annexin V^+^, PI ^+/-^), as a percentage of all events, for uninfected A549 cells treated with the listed concentration of and type of drug for 24 hours. **(E)** Same as in D, but A549 cells were infected with Ad5 for two hours before subsequent 24 hour treatment with listed drugs and apoptosis assay performed a total of 26 hpi. Statistical significance for flow cytometry data was analyzed by lognormal ordinary one-way ANOVA, and displayed as P-value >0.5 (not-significant, n.s.), * P< 0.05, ** P< 0.01 and *** P< 0.001.

**Supplementary Figure 3.**
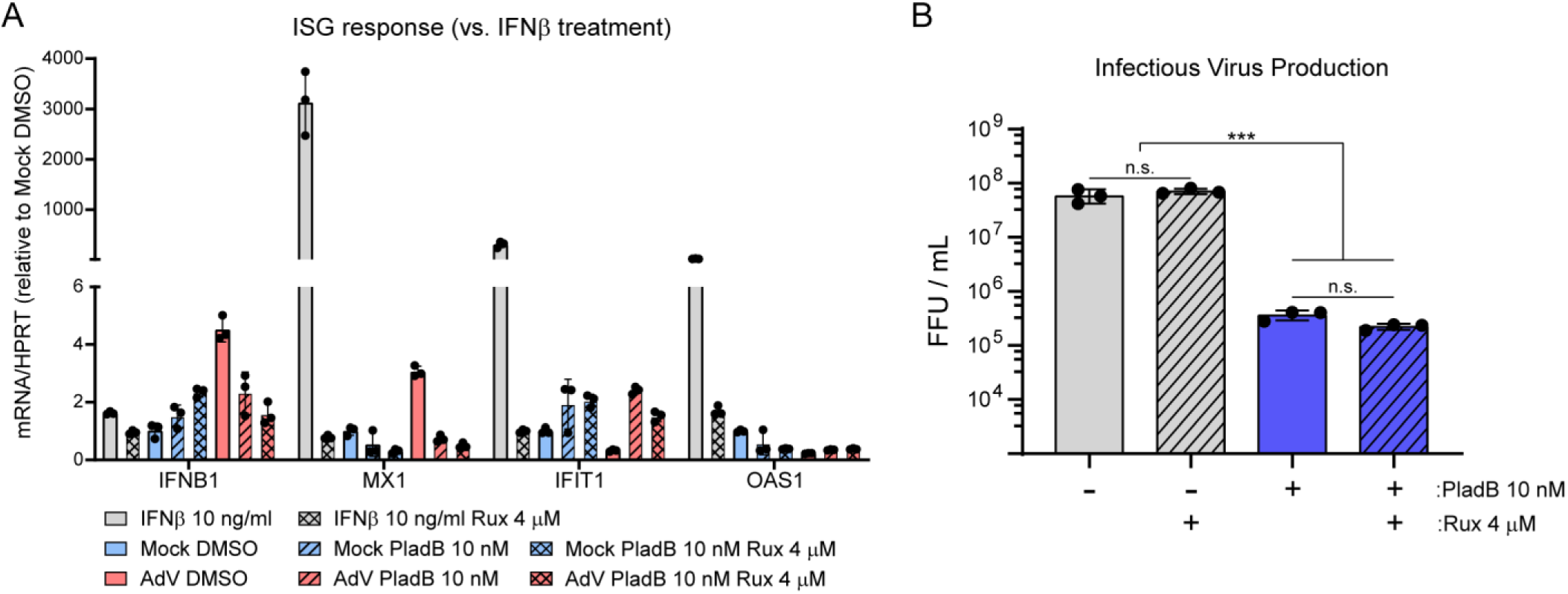
Viral specific effects of splicing inhibition are independent from an interferon stimulated response. **(A)** A549 cells were mock-infected or Ad5-infected for two hours before treatment with either 10 nM PladB alone or 10 nM PladB plus 4 μM ruxolitinib (Rux). As a control, separate wells of uninfected A549 cells were treated with 10 ng/mL recombinant Interferon Beta (IFNβ) alone or in combination with concurrent 4 μM rux treatment. Twenty-four hours post infection total RNA was harvested and subjected to qRT-PCR for interferon beta transcripts or select interferon stimulated genes. Individual transcripts were normalized to internal housekeeping control HPRT and displayed as fold over the average mock DMSO-treated cells. Bars represent the average of three biological replicates, and error bars denote standard deviation. **(B)** A549 cells were infected with fluorescent Ad5 (E3mNG) and treated with 10 nM PladB alone, 4 μM Rux alone, or 10 nM PladB plus 4 μM Rux at 2 hpi. Total cell plus supernatant was collected at 48 hpi and released viruses used for virus progeny calculation via fluorescent forming unit (FFU). Data are presented as log-scale, points show the average of three biological replicates, and error bars show standard deviation. Significance was analyzed by unpaired two-tailed *t*-test (or lognormal ordinary one-way ANOVA for infectious titer data) and displayed as P-value >0.5 (not-significant, n.s.), * P< 0.05, ** P< 0.01 and *** P< 0.001.

**Supplementary Table 1.**
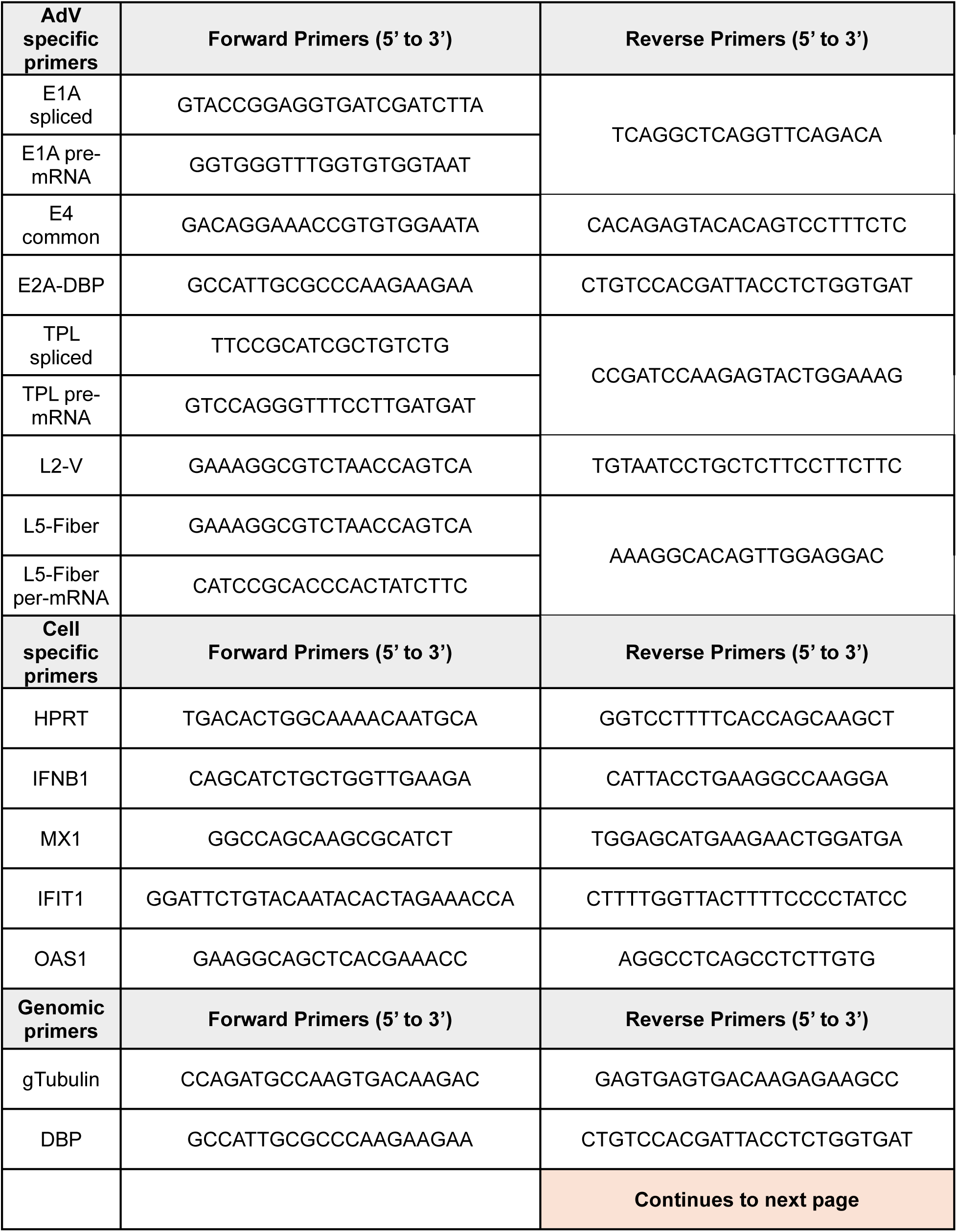
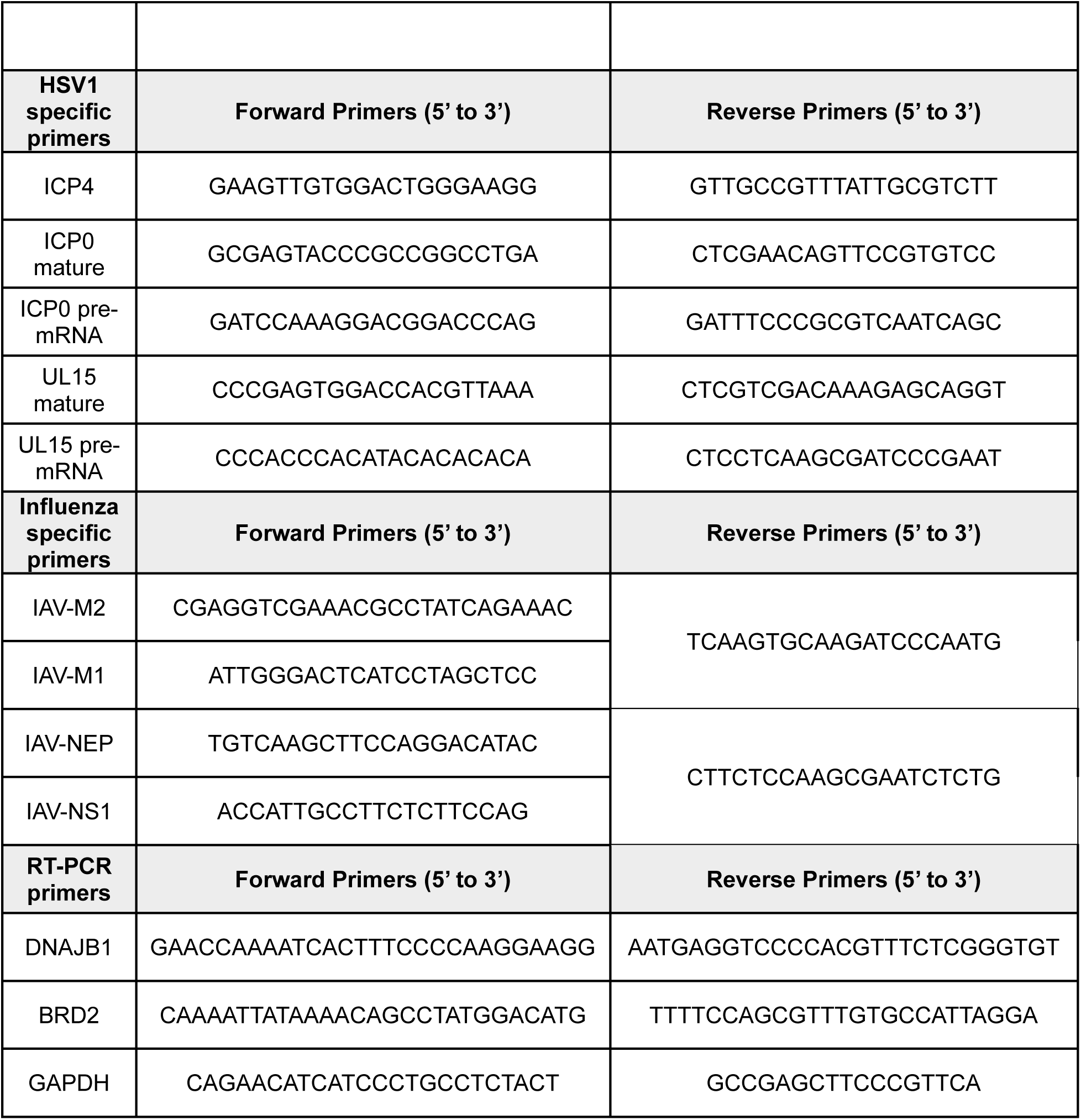
List of primers used in this study.

## Notes

### Competing Interest Statement

The authors have declared no competing interest.

